# Overcoming clinical resistance to EZH2 inhibition using rational epigenetic combination therapy

**DOI:** 10.1101/2023.02.06.527192

**Authors:** Yaniv Kazansky, Daniel Cameron, Helen Mueller, Phillip Demarest, Nadia Zaffaroni, Noemi Arrighetti, Valentina Zuco, Yasumichi Kuwahara, Romel Somwar, Marc Ladanyi, Rui Qu, Elisa De Stanchina, Filemon Dela Cruz, Andrew Kung, Mrinal Gounder, Alex Kentsis

**Author notes:** Correspondence: Alex Kentsis, MD, PhD.

## Abstract

Essential epigenetic dependencies have become evident in many cancers. Based on the functional antagonism between BAF/SWI/SNF and PRC2 in *SMARCB1*-deficient sarcomas, we and colleagues recently completed the clinical trial of the EZH2 inhibitor tazemetostat. However, the principles of tumor response to epigenetic therapy in general, and tazemetostat in particular, remain unknown. Using functional genomics of patient tumors and diverse experimental models, we sought to define molecular mechanisms of tazemetostat resistance in *SMARCB1*-deficient sarcomas and rhabdoid tumors. We found distinct classes of acquired mutations that converge on the RB1/E2F axis and decouple EZH2-dependent differentiation and cell cycle control. This allows tumor cells to escape tazemetostat-induced G1 arrest despite EZH2 inhibition, and suggests a general mechanism for effective EZH2 therapy. This also enables us to develop combination strategies to circumvent tazemetostat resistance using cell cycle bypass targeting via AURKB, and synthetic lethal targeting of PGBD5-dependent DNA damage repair via ATR. This reveals prospective biomarkers for therapy stratification, including PRICKLE1 associated with tazemetostat resistance. In all, this work offers a paradigm for rational epigenetic combination therapy suitable for immediate translation to clinical trials for epithelioid sarcomas, rhabdoid tumors, and other epigenetically dysregulated cancers.

**Significance:** Genomic studies of patient epithelioid sarcomas, rhabdoid tumors, and their cell lines identify mutations converging on a common pathway that is essential for response to EZH2 inhibition. Resistance mutations decouple drug-induced differentiation from cell cycle control. We identify complementary epigenetic combination strategies to overcome resistance and improve durability of response, supporting their investigation in clinical trials.

## Introduction

Unlike conventional cytotoxic chemotherapy, epigenetic therapy offers the ability to target cancer-specific dependencies with increased specificity and reduced toxicity. This is especially true for cancers caused by genetic mutations of transcriptional regulators, such as the chromatin remodeling BAF/SWI/SNF (Brg/Brahma-associated factors) complex that is mutated in more than 20% of human cancers (1). For example, malignant rhabdoid tumors (MRT) and epithelioid sarcomas (ES) are lethal tumors of children and young adults caused by inactivating mutations of *SMARCB1/SNF5/INI1/BAF47*, one of the core BAF/SWI/SNF complex subunits (2–4). Loss of normal BAF function can confer a dependency on the Polycomb Repressive Complex 2 (PRC2) and its methyltransferase EZH2. This dependency is thought to result from epigenetic antagonism between the two complexes, in which normal BAF activity evicts PRC2 from tumor suppressor loci (5, 6).

High *EZH2* expression in many cancer types and gain-of-function *EZH2* mutations in lymphomas (7–10) have led to the development of multiple inhibitors of EZH2 (11–13), several of which have now entered clinical trials (14). This includes clinical trials for *SMARCB1*-deficient tumors without *EZH2* mutations (15). However, despite the potential of such targeted epigenetic therapies, the principles governing their effective application remain unknown. Intrinsic and acquired resistance limits their use as monotherapies (16, 17).

A recent clinical trial led to the FDA approval of the EZH2 methyltransferase inhibitor tazemetostat (TAZ) as the first targeted therapy for *SMARCB1*-deficient epithelioid sarcomas (15). However, only 15% of patients showed objective clinical responses, with most epithelioid sarcomas being resistant to TAZ. Ongoing clinical trials in patients with rhabdoid tumors have shown similar results, with only a subset of brain atypical teratoid rhabdoid tumor (ATRT) patients exhibiting objective responses to TAZ, while extracranial rhabdoid tumors appear to be uniformly resistant (18). Recent studies have nominated the histone methyltransferase NSD1 as a regulator of TAZ susceptibility in rhabdoid tumor cells, but the clinical relevance of this mechanism is currently not known (19). In all, the outcomes for most patients with MRT and ES remain dismal, and the mechanisms of clinical response and resistance to EZH2 inhibition remain unknown. This hinders our ability to stratify treatment using prospective biomarkers to identify patients who may benefit from TAZ and to develop effective combination therapies with improved and durable benefits for patients.

Here, we define the key requirements for effective epigenetic therapy in diverse *SMARCB1*-deficient epithelioid sarcomas and rhabdoid tumors *in vivo*. Using comparative genomic analyses of clinical trial patients treated with TAZ, we identify multiple acquired mutations that cause therapy resistance. Using functional genomic approaches, we show that resistance mechanisms converge on a common RB1/E2F axis that integrates control of tumor cell division and differentiation. This organizes patient resistance mutations into a general framework, allowing us to develop rational combination therapies to circumvent TAZ resistance. Using diverse patient-derived ES and MRT cell lines and mouse xenografts, we demonstrate cell cycle bypass and synthetic lethal treatment strategies suitable for immediate translation to combination clinical trials for patients.

## Results

### Patient tumor sequencing reveals diverse resistance mutations

To identify mutations associated with clinical resistance to TAZ, we performed targeted gene sequencing of patient tumors using MSK-IMPACT, which is based on a panel of over 500 genes recurrently mutated in diverse cancer types (20). We analyzed 33 tumor specimens from 20 patients treated as part of the recent TAZ clinical trial (15), and identified somatic tumor mutations in matched pre- and post-treatment specimens (**Supplementary Tables 1 and 2**). We found distinct sets of somatic mutations in responding and non-responding tumors, with nearly all mutations, apart from *SMARCB1* loss itself, being exclusive to either TAZ-responsive or TAZ-resistant tumors (**Figure 1A, top panel**). Strikingly, we observed two tumors which initially responded to TAZ based on radiographic imaging but later developed clinical resistance (**Figure 1A, bottom panel, 1B**). Targeted sequencing of the resistant tumors revealed two newly acquired somatic mutations: heterozygous missense mutation of *EZH2* (*EZH2^Y666N^*) in the Patient 3 tumor specimen, and biallelic loss of function mutation of *RB1*, including a hemizygous deletion and a frame shift mutation (*RB1^del^*) in the remaining allele, in the Patient 15 tumor specimen. We confirmed both mutations using RNA-seq of the respective tumor specimens (**Figure S1A-B**). Since one mutation affected *EZH2* directly, and the other involved the known EZH2 target *RB1* (21), we hypothesized that both mutations were responsible for TAZ resistance in their respective patients.

**Figure 1:**
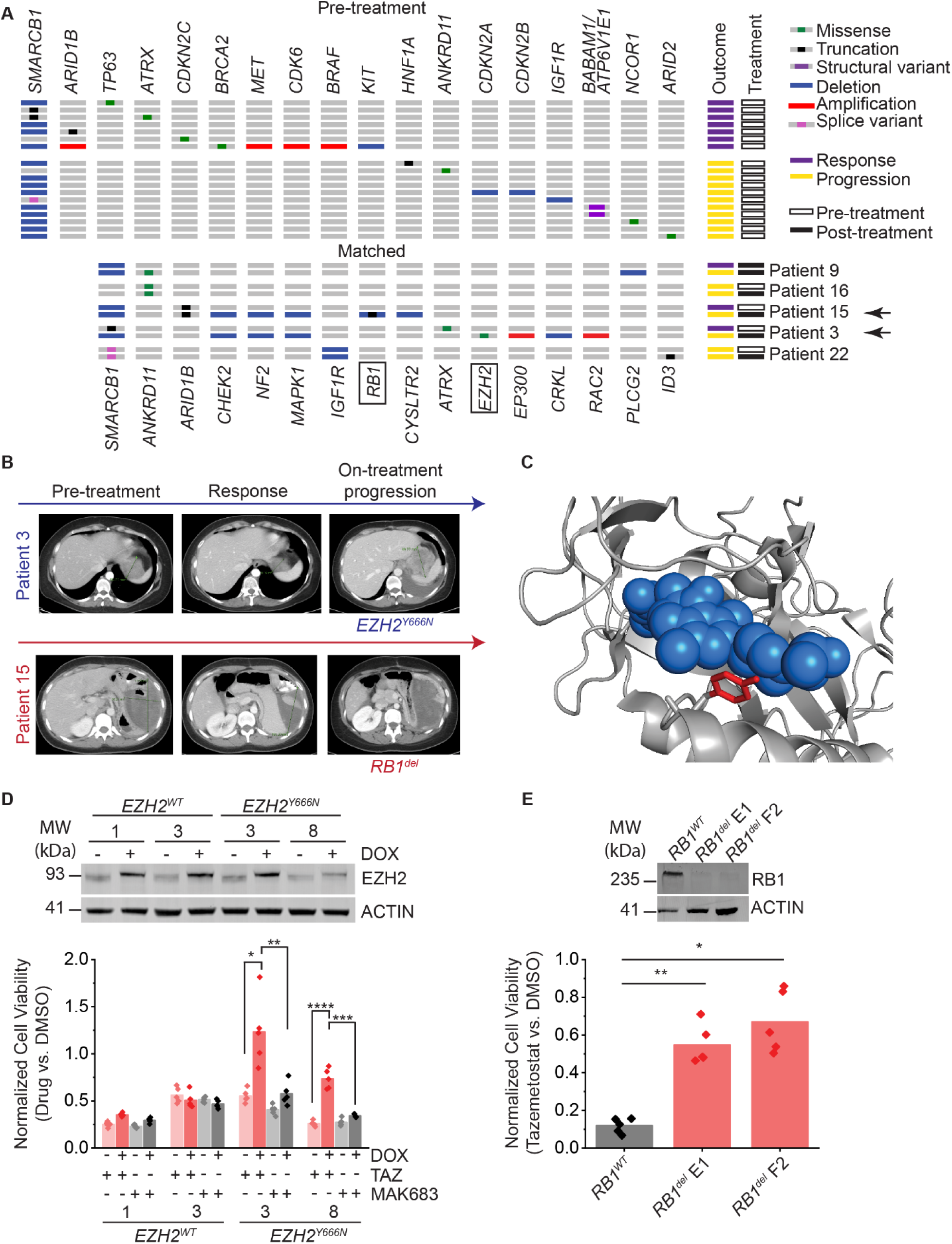
Patient tumor sequencing reveals diverse tazemetostat resistance mutations. (**A**) Abridged oncoprint of selected genes from MSK-IMPACT sequencing on patient tumor samples. Top panel: Tumor samples prior to TAZ treatment. Bottom panel: Matched samples pre- and post-TAZ or pre- and post-acquisition of resistance. (**B**) Pre- and post-treatment CT imaging of the indicated patient tumors which acquired *EZH2* and *RB1* mutations. (**C**) Atomic molecular model of the chimeric *Homo sapiens/Anolis carolinensis* EZH2 bound to pyridone-based EZH2 inhibitor I (blue), PDB: 5IJ7. Y666 is highlighted in red. (**D**) Top panel: Doxycycline-inducible *EZH2* is expressed in single-cell G401 clones at near-physiological levels after 3 days treatment with doxycycline at 1 µg/mL. Numbers indicate clone ID. Bottom panel: Cell viability measured by CellTiter-Glo after 14 days of treatment with the indicated drug at 10 µM or equivalent volume of DMSO. n=5 biological replicates per condition. \**p* = 3.5E-3, \*\**p* = 5.2E-3, \*\*\**p* = 3.8E-5, \*\*\*\**p* = 1.1E-5 by two-sided Student’s t-test. (**E**) Top panel: *RB1* knockout in two G401 clones (E1 and F2). Bottom panel: Cell viability measured by CellTiter-Glo after treatment with 10 µM tazemetostat or DMSO for 14 days. *n*=5 biological replicates per condition. \**p* = 2.8E-5, \*\**p* = 9.0E-5 by two-sided Student’s t-test.

First, we investigated the *EZH2^Y666N^* mutation. Past studies using forward genetic screens in lymphoma cell lines have identified putative resistance mutations within both the N-terminal D1 domain and the catalytic SET domain of EZH2, both of which are predicted to interact with S-adenosyl methionine (SAM)-competitive EZH2 inhibitors such as TAZ (22, 23). One such SET domain mutation previously identified in cell lines is *EZH2^Y661D^*. Y661 corresponds to Y666 in isoform 2 of *EZH2*, the isoform referred to in this study, and thus is concordant with the mutation we observed in the Patient 3 tumor (22).

Based on the atomic resolution structure of a pyridone-based EZH2 inhibitor bound to the *Anolis carolinensis* EZH2 (PDB 5IJ7) (24), we reasoned that residue Y666 in human EZH2 may form a critical part of the TAZ binding site and that its mutation can prevent TAZ from binding to the SET domain (**Figure 1C**). To test *EZH2^Y666N^*as a resistance allele, we expressed doxycycline-inducible *EZH2^Y666N^*in *SMARCB1*-deficient G401 rhabdoid tumor cells. We observed that *EZH2^Y666N^*expressing clones are resistant to TAZ as compared to cells expressing equal levels of wild-type *EZH2* by assessing both cell viability (**Figure 1D**) and cell morphology (**Figure S1C**). The resistance phenotype of *EZH2^Y666N^*-expressing cells depends on the intact catalytic activity of the SET domain, as the compound mutant combining the catalytically inactive triple mutant *EZH2^F672I,H694A,R732K^*(*EZH2^CatMut^*) with the Y666N mutation, termed *EZH2^QuadMut^*, did not confer resistance to TAZ (**Figure S1D**). This indicates that the *EZH2^Y666N^*mutation does not significantly interfere with enzymatic activity. We also observed that *EZH2^Y666N^* confers resistance to the dual EZH1/2 inhibitor valemetostat (25), consistent with putative resistance to SAM-competitive, pyridone-based EZH2 inhibitors (**Figure S1E**). This suggests that combined inhibition of EZH2 and EZH1 may not overcome this type of acquired EZH2 inhibitor resistance.

Previous studies found that lymphoma cells resistant to EZH2 inhibitors remained susceptible to the inhibition of the non-enzymatic PRC2 subunit EED (26), including those with mutations in the *EZH2* SET domain (*EZH2^C663Y^* and *EZH2^Y726F^*) and the D1 domain (23, 27). We therefore hypothesized that TAZ resistance conferred by *EZH2^Y666N^* could be overcome by PRC2 inhibitors that do not bind to EZH2. Indeed, we found that the allosteric EED inhibitor MAK683 overcomes *EZH2^Y666N^-*mediated resistance (28), demonstrating that these cells remain generally susceptible to PRC2 inhibition (**Figure 1D**). EED inhibition may thus be an effective strategy to overcome acquired TAZ resistance mutations in *EZH2*.

### *RB1* loss allows escape from cell cycle arrest despite effective EZH2 inhibition

Past work has shown that *EZH2* is a direct target of repression by RB1/E2F (21, 29). This suggests that acquired *RB1* loss may confer resistance to EZH2 inhibition by increasing EZH2 expression. To test *RB1^del^*as a TAZ resistance allele, we used CRISPR/Cas9 genome editing to generate biallelic *RB1^del^*mutations in G401 cells, as compared to the isogenic *RB1*-wild type control cells, engineered by targeting the safe harbor locus *AAVS1*. We confirmed absence of RB1 protein expression in two independent clones using Western blotting, and found that *RB1^del^* cells were indeed resistant to TAZ (**Figure 1E**). Even when the TAZ dosage is increased as high as 10 μM, this resistance persists unabated.

Despite *EZH2* being a known target gene of RB1/E2F, we were surprised to observe that *RB1^del^* G401 cells showed similar morphological changes upon TAZ treatment (**Figure S2A**) to those previously reported for TAZ-treated *RB1^WT^* G401 cells (11). To define the effects of TAZ on *RB1^del^* cells more precisely, we performed RNA-seq of isogenic G401 *RB1^del^* and wildtype *AAVS1*-control cells, treated with either 10 µM TAZ or DMSO control for 11 days, based on an established treatment regimen to model EZH2 inhibition *in vitro* (11). As predicted, we observed that *EZH2* mRNA and protein levels remained high in TAZ-treated *RB1^del^*cells, unlike in *RB1^WT^* cells (**Figure 2A-C, S2B**). However, EZH2 inhibition induced significant upregulation of hundreds of genes in both *RB1^WT^* and *RB1^del^* cells, including upregulation of known PRC2 target genes (**Figure 2A-F**).

**Figure 2:**
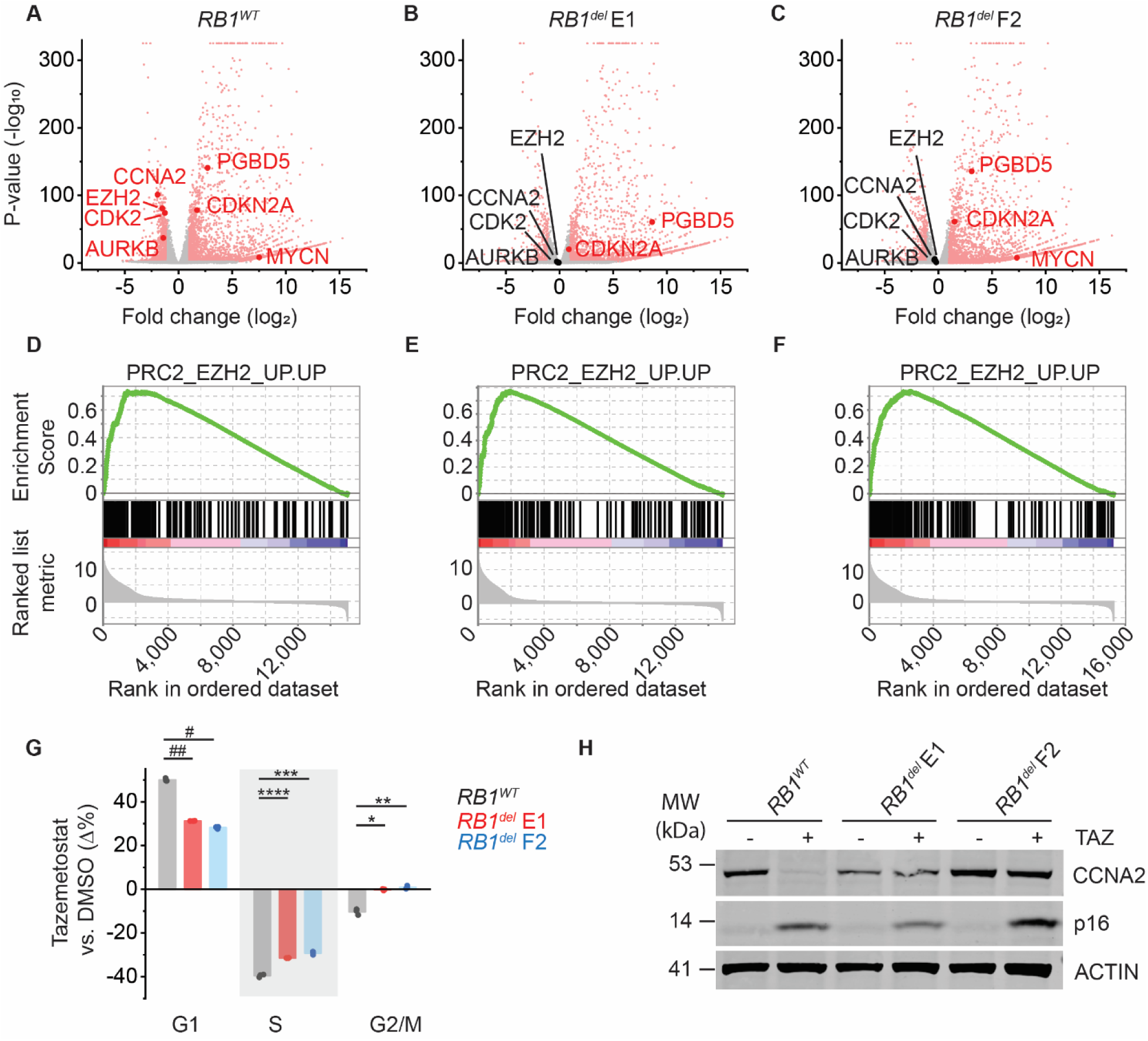
*RB1* loss allows escape from cell cycle arrest despite effective EZH2 inhibition. (**A-C**) Volcano plots of RNA-seq data from TAZ-treated G401 *RB1^WT^* (**A**), *RB1^del^ E1* (**B**), and *RB1^del^ F2* (**C**). Plots show gene expression changes of cells treated with 10 µM TAZ versus equivalent volume of DMSO for 11 days. n=3 biological replicates per condition. Dots in red show genes with expression changes of log_2_(fold change) > ±1 and *p*-value < 0.01. (**D-F**) GSEA plots showing the PRC2_EZH2_UP.UP gene set for G401 *RB1^WT^* (**D**), *RB1^del^ E1* (**E**), and *RB1^del^ F2* (**F**) cells treated in A-C. (**G**) Cell cycle analysis of the indicated G401 clone. Plot shows triplicate measurements of cells treated with 1 µM TAZ versus equivalent volume of DMSO for 11 days. Y-axis shows the percent difference of the TAZ-treated cells in each cell cycle phase compared to DMSO-treated cells. \**p=*2.4E-4, \*\**p*=2.3E-4, \*\*\**p*=5.0E-5, \*\*\*\**p*=2.7E-5, #*p*=1.4E-6, ##*p*=1.1E-6. (**H**) Western blot of the indicated G401 cell clones treated with 10 µM TAZ versus equivalent volume of DMSO for 11 days. Actin serves as loading control.

Importantly, trimethylation of the EZH2 substrate H3K27 was substantially reduced by TAZ regardless of *RB1* status (**Figure S2B**). This indicates that despite persistent EZH2 expression, EZH2 methyltransferase activity is effectively inhibited by TAZ regardless of *RB1* loss. Gene set enrichment analysis (GSEA) showed that all three clones significantly upregulated multiple gene sets upon TAZ treatment (**Supplementary Table S3**, **Figure S2C**). This included gene sets associated with mesenchymal cell differentiation such as epithelial-to-mesenchymal transition (EMT; **Figure S2D**), as well as specific markers of differentiation such as *MMP2* (**Figure S3A**), used previously as a mesenchymal marker induced by *SMARCB1* re-expression in rhabdoid tumor organoids (30). This is reminiscent of recent observations that re-expression of *SMARCB1* in G401 cells can lead to a mesenchymal chromatin state (31), and is consistent with the idea that PRC2 inhibition may allow BAF to re-activate a more developmentally normal gene expression state. While MMP2 protein did not appear to be upregulated by TAZ (**Figure S3B-C**), the induction of mesenchymal genes suggests that TAZ can induce a transcriptional differentiation program regardless of *RB1* status. Taken together, these findings indicate that *RB1* loss-induced TAZ resistance is independent of EZH2.

In addition to *EZH2*, other RB1/E2F target genes were also upregulated in TAZ-treated *RB1^del^* cells compared to TAZ-treated *RB1^WT^* control cells (**Figure S4A-B**). Given the function of RB1 in the regulation of the G1/S cell cycle checkpoint, we hypothesized that *RB1* loss could allow cells to escape TAZ-induced cell cycle arrest. Flow cytometry cell cycle analysis showed that G401 cells treated with TAZ doses as low as 1 µM arrest at the G1/S checkpoint, as reported previously (11). However, *RB1^del^*cells exhibited a significant reduction in the proportion of cells in G1 phase upon TAZ treatment (50% of TAZ-treated *RB1^WT^* cells versus 31% and 28% for *RB1^del^* E1 and F2 clones, respectively; Student’s t-test *p* = 1.4E-6 and 1.1E-6 for *RB1^WT^* versus *RB1^del^* E1 and F2, respectively), with a corresponding increase of the proportion of cells remaining in S and G2/M phases (**Figure 2G**). In agreement with these findings, we observed persistent mRNA expression of S/G2/M-phase-associated *CCNA2*, *CDK2*, and *AURKB* genes in *RB1^del^* cells upon TAZ treatment (**Figure 2A-C**), despite the upregulation of the CDK4/6 inhibitor *CDKN2A* (p16), a known PRC2 target in MRT (5, 32, 33). We confirmed persistent maintenance of the S-phase cyclin A2 (CCNA2) protein levels despite p16 upregulation using Western blotting (**Figure 2H**). Together, these results show that *RB1* loss is sufficient to evade TAZ-induced cell cycle arrest at the G1/S restriction point despite maintaining the expected global transcriptional response to EZH2 inhibition, including upregulation of cell cycle inhibitor genes.

### Intact RB1/E2F axis is required for TAZ susceptibility

The requirement for *RB1* expression in the therapeutic response to TAZ suggests that an intact RB1/E2F axis may be a general requirement for effective EZH2 inhibitor therapy. This would predict that other genetic and epigenetic perturbations to the RB1/E2F axis, beyond *RB1* loss itself, would similarly confer escape from TAZ-induced cell cycle arrest. Analysis of our TAZ clinical trial treatment cohort revealed one patient tumor specimen with primary resistance to TAZ with intact *RB1* but inactivating mutations of both *CDKN2A* and *CDKN2B* (**Figure 1A**, **Supplementary Tables 1-2**), both of which are known to inhibit CDK4/6-mediated phosphorylation of RB1. Two additional specimens had missense mutations in *ANKRD11*: One tumor with primary resistance to TAZ and another which initially responded but later progressed on treatment, at which point a newly acquired *ANKRD11* mutation was detected (**Figure 1A**-Patients 9 and 16, respectively). *ANKRD11* is a known TP53 cofactor and putative tumor suppressor that contributes to TP53-mediated expression of pan-CDK inhibitor *CDKN1A* (34–36). *CDKN1A* itself is also known to be a PRC2 target in tumors (37–39), although its role in the response to EZH2 inhibition in *SMARCB1*-deficient sarcomas is currently unknown. These results converge on the dysregulation of the RB1/E2F axis as a clinical mechanism of evasion of TAZ-induced cell cycle arrest.

We hypothesized that epithelioid and rhabdoid tumor cell lines could also harbor mutations of the RB1/E2F axis. To investigate the functional determinants of tumor cell response to TAZ, we first analyzed the response to TAZ of seven MRT and four ES cell lines in which we confirmed loss of SMARCB1 protein expression in all ES and MRT cell lines using Western blotting, as compared to SMARCB1-expressing HEK293T cells (**Figure S5A**). We classified each line as sensitive or resistant based on the area under the curve (AUC) of their TAZ dose responses (AUC > 0.3 for sensitive G401, KP-MRT-NS, TTC642, A204, TM8716, KP-MRT-RY cell lines and AUC ≤ 0.3 for resistant ES1, VAESBJ, ES2, EPI544, MP-MRT-AN cell lines; **Figure S5B**). Given that TAZ treatment requires at least 4 days for the cellular reduction of methylated EZH2 substrates and at least 7 days for apparent antiproliferative effects in the rapidly-dividing G401 cell line (11), we confirmed that the apparent TAZ susceptibilities of these MRT and ES cell lines are not correlated with their proliferation rates (Pearson’s *r* = -0.073 and *p* = 0.81; **Figure S5C**).

In agreement with somatic mutations affecting the RB1/E2F axis associated with TAZ resistance in clinical tumor specimens (**Figure 1A**), we found mutations of *CDKN2A* in 4 out of 5 TAZ-resistant cell lines, *CDKN2B* in 2 out of 5 resistant cell lines, *CDKN1A* in 1 out of 5, and *ANKRD11* in 3 out of 5, as compared to no such mutations in any TAZ-sensitive MRT and ES cell lines (**Figure 3A**, **Supplementary Table 4**). While our analysis detected a reduction in copy number of *RB1* in TAZ-sensitive KP-MRT-RY cells, manual inspection of sequencing reads within the *RB1* gene revealed lack of homozygous deletion, with presumed retention of *RB1* expression (**Figure S5D**). Thus, mutations associated with the RB1/E2F axis are associated with resistance to TAZ in *SMARCB1*-deficient cell lines and patient tumors.

**Figure 3:**
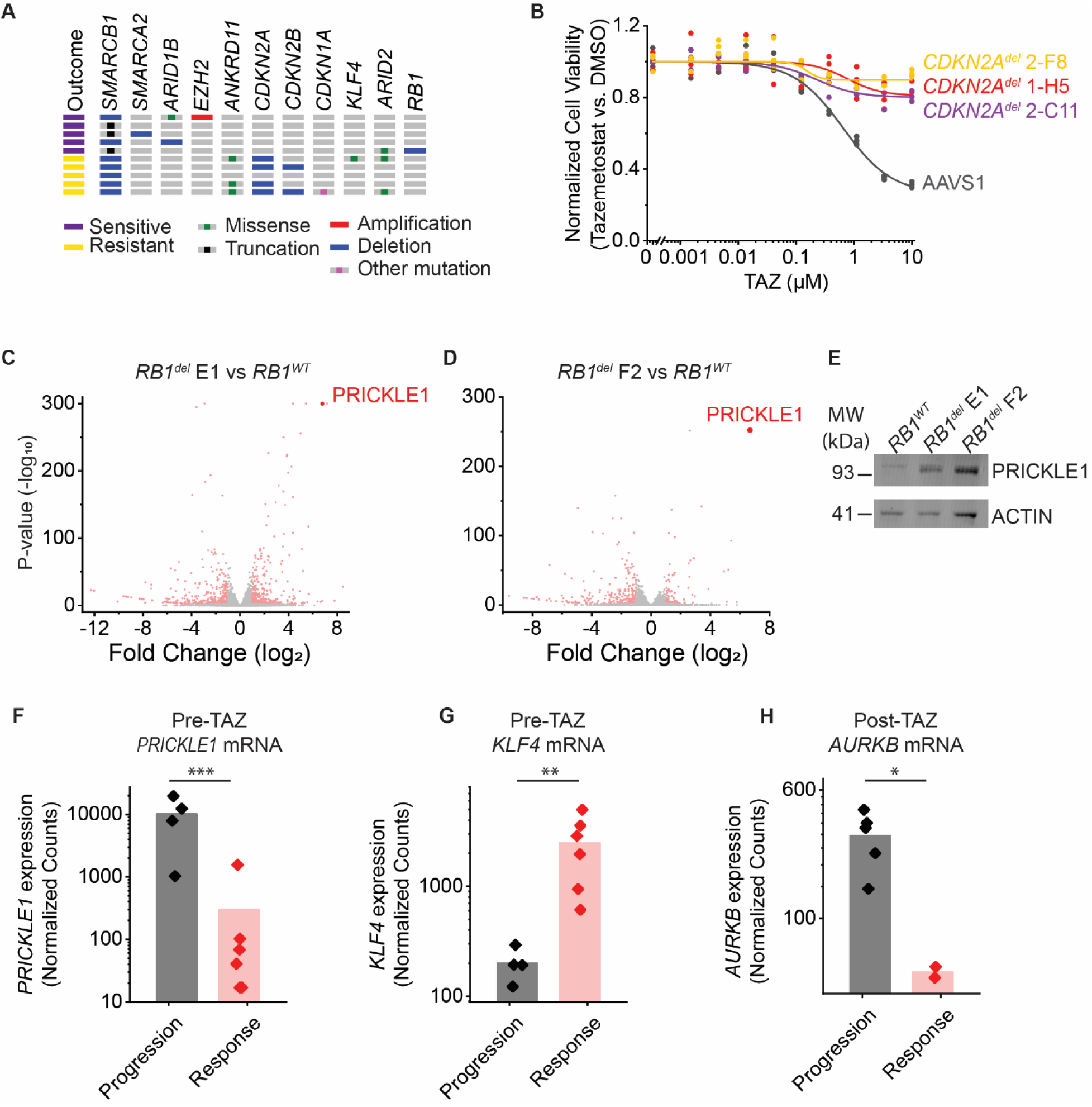
An intact RB1/E2F axis is a key requirement for response to TAZ: (**A**) Abridged oncoprint of MRT and ES cell lines. Genes included here are BAF subunits and CDK4/6/RB1/E2F axis genes. Full list of mutations is listed in Supplementary Table 4. (**B**) Indicated cell lines treated with TAZ for 11 days. (**C-D**) Volcano plots of RNA-seq data comparing DMSO-treated G401 *RB1^del^* E1 (**C**) and F2 (**D**) with *RB1^WT^* cells. (**E**) Western blot for PRICKLE1 in untreated G401 cells. (**F-G**) Read counts for *PRICKLE1* (**F**) and *KLF4* (**G**) showing counts normalized by DESeq2 for patient tumor samples collected prior to TAZ treatment. (**H**) Read counts for *AURKB* for patient tumor samples collected after TAZ treatment. \**p*=0.029, \*\**p*=0.027, *\*\*\*p*=0.013.

We confirmed that genetic mutation of *CDKN2A* correlates with lack of p16 protein induction upon TAZ treatment despite effective EZH2 inhibition by Western blotting of p16 and H3K27me3 in four TAZ-sensitive and four TAZ-resistant cell lines (**Figure S6A**). Consistent with the genomic data and observations in *RB1^del^*cells, TAZ-resistant cell lines showed no p16 induction despite complete loss of H3K27me3. Furthermore, TAZ reduced protein expression of E2F targets EZH2 and CCNA2 only in TAZ-sensitive cell lines (**Figure S6A**).

To test whether *CDKN2A* and *CDKN1A* mutations can confer resistance to TAZ, we engineered CRISPR/Cas9-mediated biallelic knockouts of *CDKN2A* or *CDKN1A* in G401 cells, generating single-cell clones with respective mutant alleles (**Figure S6B-D**). We also attempted to knock out the *ANKRD11* gene, but were unable to obtain viable biallelic mutant clones. We found that deletion of either *CDKN1A* or *CDKN2A* was sufficient to confer resistance to TAZ (**Figure S6E, 3B**). Thus, perturbations of the RB1/E2F axis beyond loss of *RB1* itself are sufficient to allow rhabdoid tumor cells to proliferate in spite of EZH2 inhibition.

We reasoned that the analysis of gene expression dynamics of TAZ-sensitive and TAZ-resistant tumor cells may be used to identify functional markers of response and resistance. Thus, we assessed the apparent transcriptional activity of the RB1/E2F axis in patient tumors using quantitative gene expression RNA-seq analysis of TAZ responding and non-responding tumors biopsied before and after TAZ treatment (**Supplementary Table S1**). Pre-treatment TAZ-resistant tumors exhibited significant enrichment of multiple Gene Ontology (GO) terms associated with the cell cycle and in particular with the S and G2/M phases (**Figure S7A**). Similarly, post-treatment tumors that progressed on TAZ showed increased gene expression of GO terms associated with mitosis, as compared to TAZ-responsive tumors (**Figure S7B**). Indeed, TAZ-resistant tumors exhibited consistently higher expression of S/G2/M-phase-associated genes prior to treatment (**Figure S7C-D**). These findings suggest that in addition to the mutations affecting the RB1/E2F axis associated with TAZ resistance, additional mutations not captured by MSK-IMPACT targeted gene sequencing and/or epigenetic dysregulation, such as putative silencing of tumor suppressor genes like *CDKN1A* or *CDKN2A*, may contribute to the decoupling of RB1/E2F-mediated proliferation and PRC2-regulated differentiation, and consequent TAZ resistance.

Since TAZ-resistant MRT and ES cell lines and patient tumors show distinct mutations and gene expression changes that converge on the RB1/E2F axis, we inquired whether these perturbations would similarly converge on common prognostic biomarkers of TAZ resistance. Comparative gene expression analysis of untreated *RB1^del^*cells versus *RB1^WT^* G401 cells showed a small set of consistently and significantly up- and down-regulated genes in two independent clones (**Figure S8A**). The most substantially and significantly upregulated gene associated with *RB1* loss was *PRICKLE1* (**Figure 3C-D**), which we confirmed to be overexpressed at the protein level in both *RB1^del^* clones using Western blotting (**Figure 3E**).

We also found that *PRICKLE1* was among the most differentially expressed genes between 10 pre-treatment patient tumors with response (6) and resistance (4) to TAZ, with *PRICKLE1* expression being higher in TAZ-resistant tumors (mean normalized read counts = 10,237 and 300 for resistant and responsive tumors, respectively; Student’s t-test *p* = 0.013; **Figure 3F, S8B**). *PRICKLE1* can control planar cell polarity (PCP), a key cell differentiation pathway, and has previously been implicated as a prognostic biomarker of poor prognosis in breast cancer (40, 41), acute myeloid leukemia (42), and gastric cancer (43, 44). Several other genes encoding PCP pathway factors were also upregulated in TAZ-resistant tumors compared to TAZ-sensitive tumors (**Figure S8B**). We also examined differentially expressed genes in TAZ-sensitive tumors as potential markers of sensitivity. These included the transcription factor *KLF4* which can control the G1/S transition by regulating *CDKN1A* expression (45, 46) (mean normalized read counts = 200 and 2,489 for resistant and responsive tumors, respectively; Student’s-test *p* = 0.03; **Figure S8C, 3G**). Thus, PRICKLE1 and possibly additional factors controlling PCP and integration of RB1/E2F cell cycle and differentiation are potential prognostic pre-treatment biomarkers to identify clinical TAZ resistance and susceptibility of *SMARCB1*-deficient tumors.

### Synthetic lethal and cell cycle bypass epigenetic combination strategies overcome tazemetostat resistance

Given that *RB1^del^* cells are able to bypass cell cycle arrest at the G1/S checkpoint, we reasoned that inhibiting cell cycle kinases downstream of this checkpoint could overcome TAZ resistance. We focused on the cell cycle kinases *CDK2* and *AURKB*, which are downregulated by EZH2 inhibition in TAZ-sensitive cells but persistently expressed in TAZ-resistant cells (**Figure 2A-C**), as potential cell cycle bypass targets. We found that the CDK2 inhibitor seleciclib (47), the mitotic kinase Aurora A inhibitor alisertib (48), and the Aurora B inhibitor barasertib (49) were able to overcome TAZ resistance in *RB1^del^* G401 cells (**Figure 4A, Figure S9A-B**). Consistent with the function of CDK4/6 kinases upstream of RB1/E2F, sensitivity to CDK4/6 inhibitors palbociclib and abemaciclib was reduced by *RB1^del^* mutation (**Figure S9C-D**). In support of the cell cycle bypass TAZ combination strategy, we observed that patient tumors which progressed on TAZ showed higher expression of *AURKB* mRNA as compared to those that responded (**Figure 3H**). Combined with the high sensitivity of G401 cells to barasertib (**Figure 4A**; half-maximal effective concentration of 6.5 ± 0.5 nM, 5.5 ± 0.6 nM, and 5.9 ± 0.9 nM for *RB1^WT^*, *RB1^del^* E1, and F2 clones, respectively), these findings suggest that the cell cycle bypass strategy may be used to overcome TAZ resistance.

**Figure 4:**
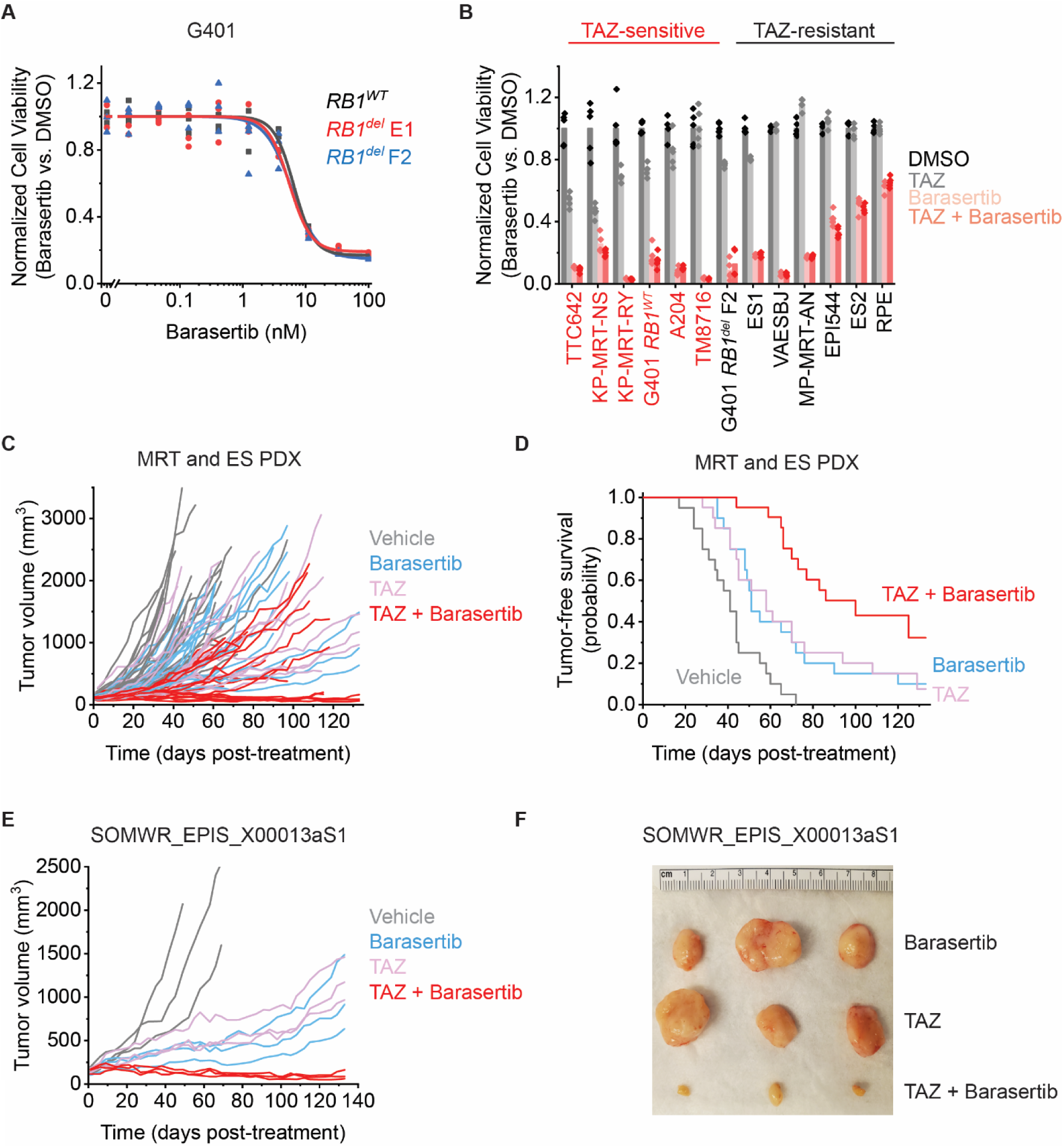
Cell cycle bypass combination strategy using AURKB inhibition overcomes TAZ resistance and improves response: (**A**) G401 cells treated with barasertib for 6 days (**A**). Panel of MRT and ES cell lines ordered left to right by decreasing response to TAZ monotherapy. Cells were treated with the indicated monotherapy or combination for 11 days. Drug concentrations used were: TAZ: 200 nM, barasertib: 8 nM. (**C**) Tumor growth curves showing volumes calculated from caliper measurements for 5 mouse PDXs treated with the indicated drug regimen. n = 20 mice for vehicle and barasertib-treated groups, n = 21 for TAZ and TAZ + barasertib-treated groups. Vardi *U*-test *p* = 4.0E-4 and 2.0E-4 for combination vs. barasertib or TAZ, respectively. (**D**) Kaplan-Meier curves showing tumor-free survival (defined as tumor volume ≤ 1,000 mm^3^) for the PDXs in panel C. Mean survival is 65 days (95% CI: 52-78 days) for barasertib, 68 days (95% CI: 53-82 days) for TAZ, 98 days (95% CI: 84-112 days) for the combination. Log-rank test *p* = 3.3E-3 and 7.4E-3 for combination vs. barasertib or TAZ, respectively. (**E**) Tumor growth curves for the subset of mouse tumors in panel C from the SOMWR_EPIS_X00013aS1 PDX model. n = 3 mice per treatment group. Vardi *U*-test *p* = 0.10 and 9.4E-2 for combination vs. barasertib or TAZ, respectively. (**F**) Tumors from panel E harvested on Day 135 of treatment.

We therefore asked whether the combination of TAZ and barasertib would have activity against both TAZ-responsive and TAZ-resistant *SMARCB1*-deficient MRT and ES cell lines, using RPE cells as a *SMARCB1*-proficient control. The effects of the combination of TAZ and barasertib on cell viability did not substantially exceed the effect of barasertib alone at the doses tested (**Figure 4B**; 200 nM TAZ, 8 nM barasertib). However, nearly all cell lines tested, including those resistant to TAZ monotherapy, showed substantial susceptibility to barasertib, with RPE cells displaying the lowest sensitivity (**Figure 4B**). Cell cycle analysis on G401 cells treated with this combination showed that the combined TAZ and barasertib treatment caused a greater degree of cell cycle arrest and reduced entry into S phase than either drug alone, as measured by EdU incorporation (**Figure S10A**). This was the case in both *RB1^WT^* and *RB1^del^* cells, consistent with the prediction that cell cycle inhibition downstream of the G1/S checkpoint would cause cell cycle arrest even in cells with *RB1* loss (**Figure S8A**). Furthermore, the S phase reduction was not due to apoptosis, as measured by Western blot and immunofluorescence analysis of Caspase 3 cleavage (**Figure S10B-C**). Finally, this effect was associated with the induction of large multi-nucleated cells (**Figure S10C**), consistent with observations of postmitotic endoreduplication upon AURKB inhibition (50, 51).

To test the effects of combined TAZ and barasertib therapy *in vivo*, we treated a panel of five patient-derived rhabdoid tumor and epithelioid sarcoma xenografts (PDX) engrafted in immunodeficient mice (**Supplementary Table S5**). In contrast to the modest reduction of tumor growth and extension of survival of mice with tumors <1,000 mm^3^ with TAZ or barasertib treatment alone, PDX mice treated with the combination of TAZ and barasertib showed significant reductions in tumor growth (Vardi *U*-test *p* = 4.0E-4 and 2.0E-4 for combination versus barasertib or TAZ, respectively; **Figure 4C**) (52). In two of the PDX models, this combination led to tumor regressions (**Figures 4E-F & S10D**). Consistent with this benefit, the combination was also found to significantly increase mean tumor-free animal survival from 65 days (95% confidence interval (CI) = 52-78 days) for barasertib and 68 days (95% CI = 53-82 days) for TAZ to 98 days (95% CI = 84-112 days) for the combination (log-rank test *p* = 3.3E-3 and 7.4E-3 for combination versus barasertib or TAZ, respectively; **Figure 4D**). These results indicate that the combination of TAZ with a downstream cell cycle inhibitor such as barasertib can improve response and overcome resistance to TAZ in diverse rhabdoid tumors and epithelioid sarcomas *in vivo*.

Our studies with MRT and ES cell lines showed that p16 induction, rather than inhibition of EZH2 and H3K27me3 loss, correlates with TAZ response. To determine whether this relationship held *in vivo*, we measured H3K27me3 and p16 levels in three PDX models for which sufficient mice were engrafted and treated. One model, HYMAD_EPIS_X0004aS1, did not respond to TAZ monotherapy (log-rank test *p* = 0.63), while the other two, SOMWR_EPIS_X00013aS1 and KUNGA_MRT_X0002aS1 showed significant anti-tumor activity of the TAZ alone (log-rank test *p* = 7.1E-6 and 8.0E-3 respectively; **Figure S11A**). While all three models showed reductions of H3K27me3 upon TAZ treatment, only TAZ-sensitive tumors showed upregulation of p16 and reduction of the cell proliferation marker Ki67 (**Figure S11B-C**). In addition, TAZ did not appear to induce apoptosis *in vivo* as evidenced by the absence of cleaved caspase 3 staining (**Figure S11B-C**), consistent with the results *in vitro* (**Figure S10B-C**).

In addition to the distinct cell cycle dynamics of TAZ resistance, we observed that TAZ treatment also caused significant increase in the expression of the *PiggyBac transposable element derived 5* (*PGBD5)* regardless of *RB1* mutation (**Figure 2A-C**). *PGBD5* is a transposase-derived gene with retained nuclease activity in human cells (53–55), which requires active end-joining DNA repair, including the DNA damage repair signaling kinase ATR (56). In rhabdoid tumors and medulloblastomas, PGBD5 has been implicated as a developmental mutator and effector of somatic deletions of *SMARCB1* itself, and has been found to induce double-strand DNA (dsDNA) breaks and DNA rearrangements (53, 56–58).

Since *PGBD5* is both necessary and sufficient to confer a dependency on ATR kinase signaling in tumor cells (56), we reasoned that TAZ-induced upregulation of *PGBD5* expression may potentiate the synthetic lethal dependency between PGBD5 and ATR-dependent DNA damage repair. To test this idea, we used the ATR-selective kinase inhibitor elimusertib, which is currently undergoing clinical trials in patients with solid tumors, including patients with PGBD5-expressing tumors such as MRT and ES (Clinical Trials Identifier NCT05071209). We found that elimusertib exhibited low-nanomolar potency against *RB1^WT^* and *RB1^del^* G401 cells *in vitro* (half-maximal effective concentration of 18 ± 1.6 nM, 19 ± 3.8 nM, and 27 ± 3.2 nM for *RB1^WT^*, *RB1^del^* E1 and F2 clones, respectively; **Figure 5A**). We also found that the combination of TAZ and elimusertib exerted greater antitumor effects than either drug alone against diverse MRT and ES cell lines (**Figure 5B**), exhibiting synergy in a subset of the cell lines (**Figure S12A-B**). To determine whether the synergistic combination of elimusertib and TAZ antitumor effects were due to increased DNA damage, we used confocal immunofluorescence microscopy to quantify γH2AX, as a specific marker of DNA damage (59). In agreement with prior studies (56), untreated G401 cells showed measurable γH2AX associated with baseline *PGBD5* expression (**Figure 5D**). Consistent with TAZ-mediated induction of *PGBD5* expression (**Figure 2A-C**), we found that TAZ treatment alone significantly increased nuclear γH2AX (mean normalized level = 0.42 versus 0.29 for TAZ and DMSO, respectively; *t*-test *p* = 9.3E-3; **Figures 5C-D, S12C**), and the combination of TAZ and elimusertib induced additional increases in γH2AX than either drug alone (mean normalized levels = 1.8 for the combination versus 0.42 and 0.85 for TAZ and elimusertib, respectively; *t*-test *p* = 6.2E-14 and 1.5E-29 for combination versus elimusertib and TAZ, respectively; **Figure 5D, S12C**).

**Figure 5:**
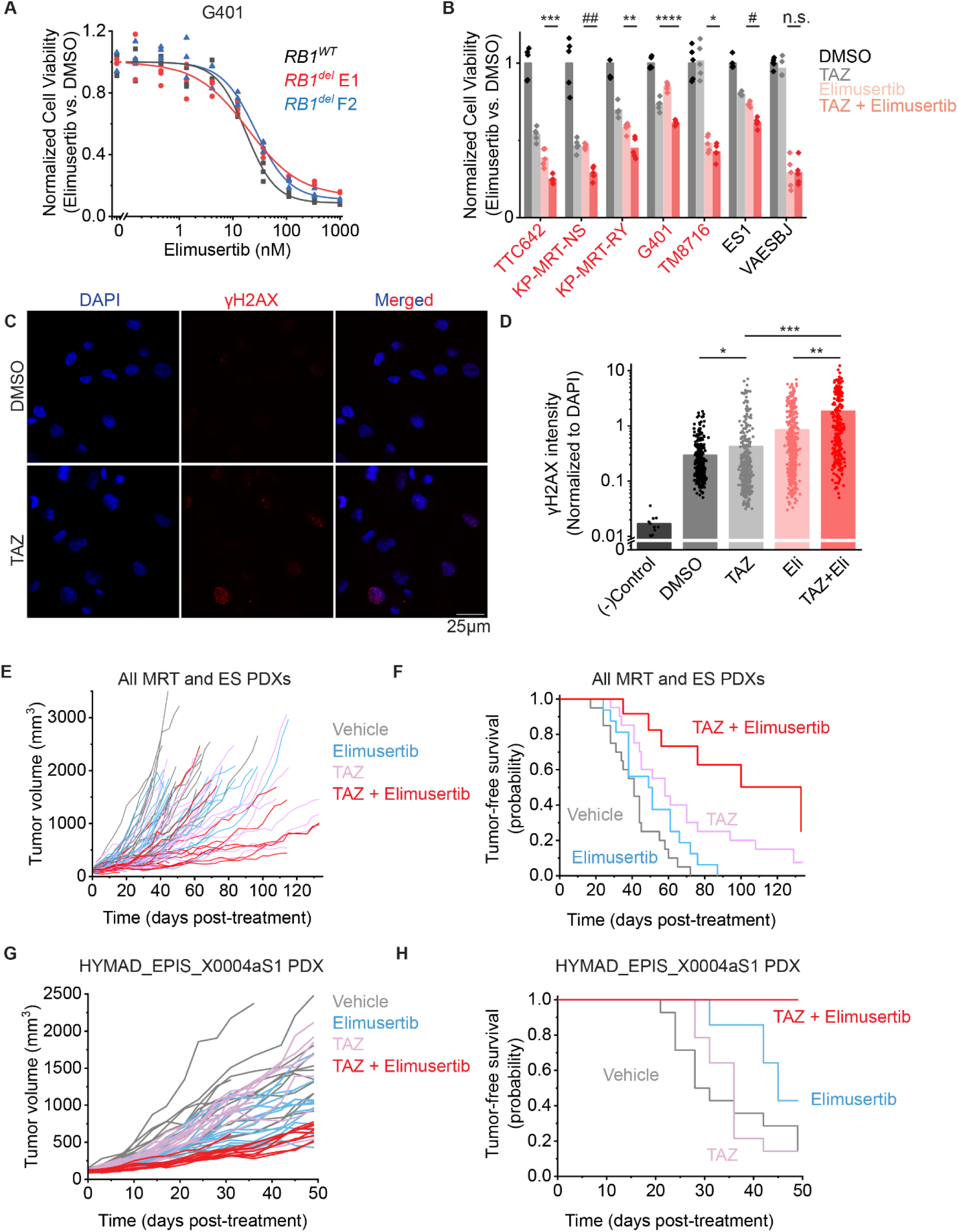
Synthetic lethal combination strategy using ATR inhibition overcomes TAZ resistance and improves response: (**A**) G401 cells treated with elimusertib for 4 days. (**B**) Panel of MRT and ES cell lines ordered left to right by decreasing response to TAZ monotherapy. Cells were treated with the indicated monotherapy or combination for 11 days. Drug concentrations used were: TAZ: 200 nM, elimusertib: 8 nM. We selected an elimusertib dose below its monotherapy IC_50_ (for G401 cells) in order to visualize any additive effects upon combination with TAZ. \**p* = 0.14, \*\**p* = 3.3E-3, \*\*\**p* = 1.0E-3, \*\*\**p* = 2.6E-4, #*p* = 6.1E-5, ##*p* = 3.0E-529 by two-sided Student’s t-test. All comparisons refer to TAZ + elimusertib vs. elimusertib conditions, except for G401 cells, in which comparison is for TAZ + elimusertib vs. TAZ. n = 5 replicates per condition. (**C**) Representative images of G401 cells treated with the indicated treatment for 7 days. Elimusertib was added on Day 5. Doses used were 500 mM TAZ and 100 nM elimusertib. (**D**) Quantification of γH2AX fluorescence relative to DAPI fluorescence. \**p* = 9.3E-3, \*\**p* = 6.2E-14, \*\*\**p* = 1.5E-29 by two-sided Student’s t-test. n = 332 nuclei for DMSO, 404 for TAZ, 400 for elimusertib, 257 for TAZ + elimusertib. (**E**) Tumor growth curves for 5 mouse PDXs treated with the indicated drug regimen. n = 20 mice for vehicle and elimusertib-treated groups, n = 21 for TAZ and TAZ + elimusertib-treated groups. Vardi *U*-test *p* = 2.3E-2 and 0.20 for combination vs. elimusertib or TAZ, respectively. (**F**) Kaplan-Meier curves showing tumor-free survival (defined as tumor volume ≤ 1,000 mm^3^) for the PDXs in panel C. Mean survival is 51 days (95% CI: 42-60 days) for elimusertib, 68 days (95% CI: 53-82 days) for TAZ to 100 days (95% CI: 74-124 days) for the combination. Log-rank test *p* = 5.8E-4 and 3.8E-2 for combination versus elimusertib or TAZ, respectively. (**G**) Tumor growth curves for the HYMAD_EPIS_X0004aS1 PDX model treated with the indicated drug regimen. Vardi *U*-test *p* = 2.0E-4 for combination versus elimusertib or TAZ. n = 14 mice per treatment group. (**H**) Kaplan-Meier curves showing tumor-free survival (defined as tumor volume ≤ 1,000 mm^3^) for the PDXs in panel E. Log-rank test *p* = 6.2E-3 and 6.3E-5 for combination versus elimusertib or TAZ, respectively.

To determine whether TAZ-mediated induction of DNA damage, associated with increased expression of *PGBD5*, is indeed PGBD5-dependent, we engineered short hairpin RNA-mediated knockdown of *PGBD5* in G401 cells using two specific shRNA constructs (shPGBD5), as compared to a non-targeting control GFP targeting shRNA (shGFP; **Figure S13A**). We found that while the combination of TAZ and elimusertib induced DNA damage in shPGBD5 cells, this effect was significantly reduced compared to control shGFP cells (Student’s t-test, *p* = 3.9E-8 and 4.4E-8 for shGFP versus shPGBD5-1 and shPGBD5-3, respectively; **Figure S13B**). This was the case both when examining all nuclei with γH2AX (**Figure S13B**), as well as analyzing nuclei with punctate γH2AX staining (**Figure S13C**), which have more localized DNA damage versus those with pan-nuclear γH2AX staining (**Figure S13D**), which are cells with genome-wide unrepaired DNA damage including cells undergoing apoptosis. Thus, the induction of DNA damage by the combination of TAZ and elimusertib is PGBD5-dependent, as explained by the synthetic lethal dependency between PGBD5 and end-joining DNA repair and the TAZ-mediated increase in PGBD5 expression and its activity as a DNA nuclease that induces DNA damage.

This synthetic lethal targeting strategy is specific, because the combination of TAZ with the SRA737 inhibitor of another DNA damage repair signaling kinase CHK1 showed no increased activity as compared to either drug alone (**Figure S14A-B**). Indeed, TAZ treatment did not induce apparent replication stress, as measured by RPA phosphorylation (**Figure S14B**), which was also not potentiated by the combined CHK1 inhibition with SRA737, despite effective suppression of CHK1 auto-phosphorylation (**Figure S14B**). Unlike T-ALL, where EZH2 suppression induces MYCN protein expression and replication stress (60), TAZ treatment of G401 rhabdoid tumor cells failed to increase MYCN protein levels (**Figure S14C**), in spite of significant upregulation of *MYCN* mRNA (**Figure 2A-C**).

Encouraged by the potent and specific antitumor activity of synthetic lethal combination TAZ therapy *in vitro*, we tested the antitumor activity of elimusertib and TAZ combination using a diverse cohort of MRT and ES PDX mice *in vivo* (**Supplementary Table S5**). The combination of TAZ and elimusertib exceeded the effect of treatment with either drug alone when assessed by tumor measurements (Vardi *U*-test *p* = 2.33E-2 and 0.20 for combination versus elimusertib or TAZ, respectively; **Figure 5E**) and significantly extended tumor-free survival from 51 days (95% CI = 42-60 days) for elimusertib and 68 days (95% CI = 53-82 days) for TAZ to 100 days (95% CI = 76-124 days) for the combination (log-rank test *p* = 5.8E-4 and 3.8E-2 for combination versus elimusertib or TAZ, respectively; **Figure 5F**). This was most pronounced for the HYMAD_EPIS_X0004aS1 and SOMWR_EPIS_X00013aS1 PDX models (**Figures 5G-H & S15A-D**), despite the former exhibiting a relatively poor response to TAZ monotherapy, when assessed by tumor growth measurements (Vardi *U*-test *p* = 2.0E-4 for combination versus elimusertib or TAZ for HYMAD_EPIS_X0003aS1; **Figure 5G**; *p* = 1.0E-3 and 6.0E-2 for combination versus elimusertib and TAZ, respectively for SOMWR_EPIS_X00013aS1; **Figure S15C**) and tumor-free survival (log-rank test *p* = 6.2E-3 and 6.3E-5 for combination versus elimusertib or TAZ, respectively for HYMAD_EPIS_X0003aS1; **Figure 5H**; *p* = 9.5E-5 and 6.0E-2 for combination versus elimusertib or TAZ, respectively for SOMWR_EPIS_X00013aS1; **Figure S15D**). Thus, the combination of EZH2 and ATR inhibition constitutes a synthetic lethal rational combination strategy to improve TAZ clinical response and overcome resistance.

Finally, we assessed the impact of TAZ treatment on immune-related gene sets, as recent studies have reported that EZH2 inhibition can increase tumor immunogenicity (61–63). We found that TAZ treatment upregulated several immune-related gene sets in both *RB1^WT^*and *RB1^del^* G401 cells, with 8 of the top 25 upregulated gene sets in all cell clones involving the immune response (**Supplementary Table S3, Figure S16A-C**). We also found that post-treatment patient tumor samples which responded to TAZ expressed higher levels of genes associated with immune-related GO terms (**Figure S16D**). Immune genes upregulated by TAZ in G401 cells included both MHC-I and MHC-II genes as well as their respective regulators NLRC5 and CIITA (**Figure S16E-F**), interferon-γ and its receptor (**Figure S16G**), and *CD274* encoding PD-L1 (**Figure S16G**). Similarly, we observed TAZ induction of multiple endogenous retroelements (**Figure S16H-I**, **Supplementary Table S6**), and the dsRNA sensor *IFIH1/MDA5* (**Figure S16G**). Further investigation of EZH2-mediated and TAZ induced antigen presentation and immune responses of *SMARCB1*-deficient sarcomas is warranted, including the investigation of the combination of immune checkpoint blockade with epigenetic TAZ combination therapies.

## Discussion

What defines effective epigenetic EZH2 inhibition therapy for *SMARCB1*-deficient epithelioid sarcomas and rhabdoid tumors? Our studies of ES patients treated with TAZ clinically demonstrate that effective inhibition of PRC2 enzymatic activity is necessary but not sufficient for durable antitumor effects. Based on clinical genomics and transcriptomics, combined with functional genetic studies of more than 15 diverse MRT and ES cell lines and patient-derived tumors *in vitro* and *in vivo*, we propose a general molecular model for effective epigenetic TAZ therapy (**Figure 6**). This model places validated *RB1* and *EZH2* TAZ resistance alleles within the context of a molecular sequence of events required for clinical TAZ response. This model also explains additional mutations associated with TAZ resistance based on their perturbation of each stage of this sequence and provides candidate prognostic biomarkers and therapeutic combination strategies. We discuss the evidence for this model below, and summarize its novel predictions and implications.

**Figure 6:**
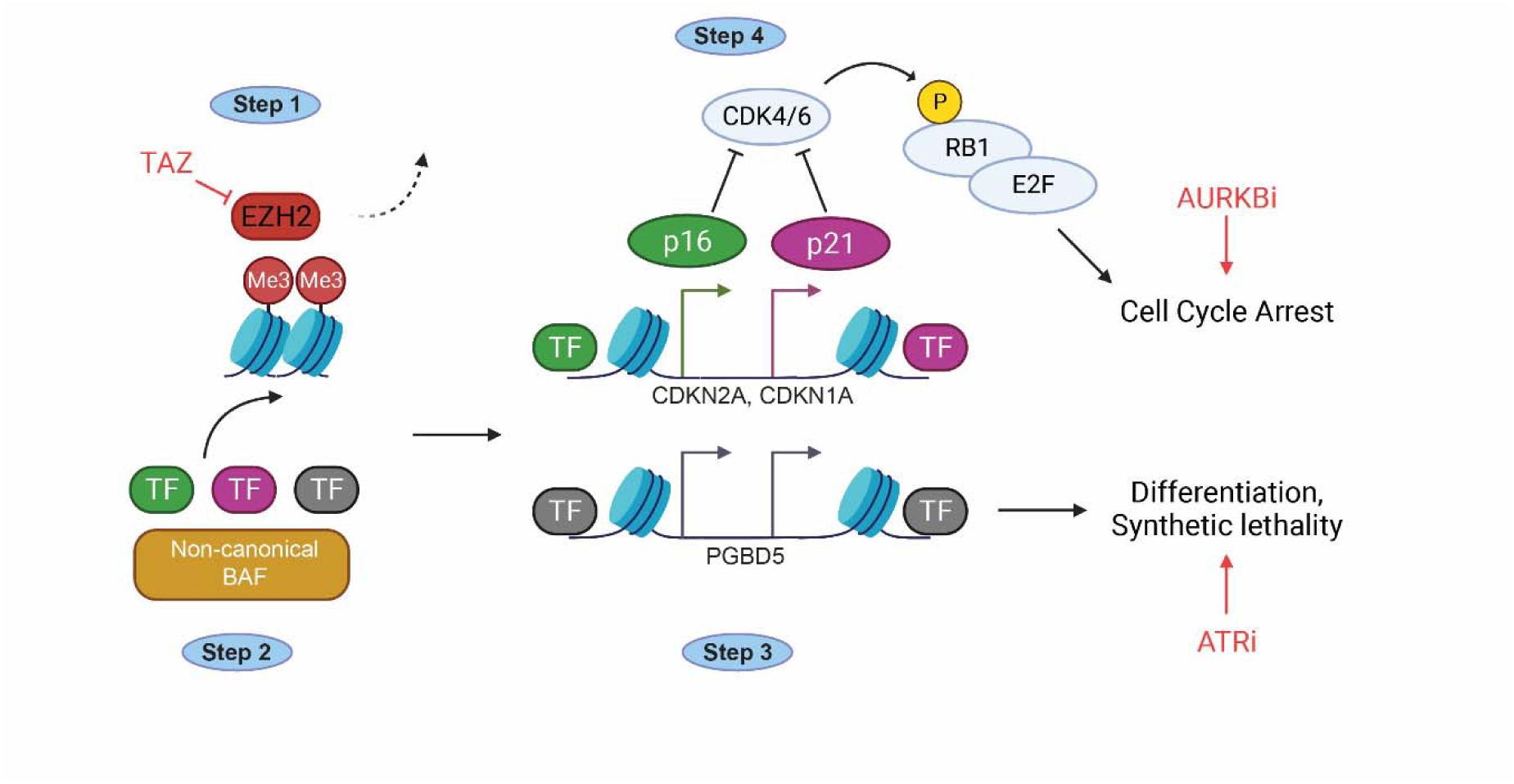
Mechanistic schematic for the response of BAF-deficient tumors to effective EZH2 therapy: **Step 1:** TAZ inhibits histone methylation activity of PRC2. **Step 2:** Activating chromatin-bound complexes, such as ncBAF bind to tumor suppressor loci. **Step 3:** Tumor suppressor loci are activated by their transcription factors and their coactivators. **Step 4:** Tumor suppressors inhibit cell cycle progression through their downstream effectors, such as RB1/E2F.

First, TAZ must be able to bind and enzymatically inhibit the EZH2 SET domain (**Figure 6; Step 1**). This inhibition can be blocked by gatekeeper mutations of the EZH2 drug binding site, as observed in lymphoma cell lines (22), and demonstrated for the first time here in an epithelioid sarcoma with clinically acquired *EZH2^Y666N^* mutation. Such resistance mutations can be overcome by targeting EED, a non-enzymatic PRC2 subunit.

For effective epigenetic TAZ therapy, chromatin remodeling complexes must act on tumor suppressor loci that were aberrantly repressed by PRC2 (**Figure 6; Step 2**). The canonical BAF complex is thought to oppose the activity of the PRC, associated with its chromatin eviction (5, 6). However, the precise mechanism of eviction of TAZ-inhibited PRC2 in *SMARCB1*-deleted tumors is not fully defined. This may involve TAZ-induced remodeling of BAF and/or PRC2. For example, a recent study of *SMARCA4*-deficient cell lines found that upregulation of the expression of BAF helicase *SMARCA2*, which is under PRC2 control in these cells, is necessary for response to EZH2 inhibition (64). In our study, we observed *RB1*-independent upregulation of the expression of BAF subunits *SMARCA2* and *DPF3* upon TAZ treatment in G401 cells (**Figure S17A**). This suggests that TAZ may impact BAF complex assembly as part of its therapeutic mechanism in *SMARCB1*-deficient tumor cells. However, this also indicates that *SMARCA2* re-expression upon EZH2 inhibition is not sufficient to induce cell cycle arrest in the absence of *RB1* expression. After assessing TAZ-induced gene expression changes in PRC2 subunits, we also observed *RB1*-independent TAZ-induced upregulation of PRC2 subunit *JARID2*, and *RB1*-dependent downregulation of *PHF19*, suggesting that PRC2 composition itself may be affected by TAZ treatment (**Figure S10A**). Additionally, it is unknown whether PRC2 eviction in TAZ-treated cells requires a specific form of the BAF complex, such as the non-canonical *SMARCB1*-deficient ncBAF or GBAF complex described previously (65, 66).

If the activity of specific BAF subtypes is indeed needed to evict PRC2 from chromatin upon EZH2 inhibition, then genetic perturbation of specific BAF subunits may impact tumor response to TAZ. For example, in our patient cohort, we observed one TAZ-sensitive tumor with a truncation in the canonical BAF-specific subunit *ARID1B*, while a TAZ-resistant tumor had a missense mutation in PBAF-specific subunit *ARID2* (**Figure 1A**). We also found *ARID1B* to be mutated in 2 out of 5 TAZ-responsive cell lines (0 out of 5 TAZ-sensitive cell lines) and *ARID2* to be mutated in 2 out of 5 TAZ-resistant cell lines, though also in 1 out of 5 TAZ-sensitive lines (**Figure 3B**). It is not known whether these mutations affect tumor response to TAZ, and further work will be needed to elucidate the specific mechanism of chromatin de-repression in TAZ-treated tumor cells, as well as the effects of TAZ on BAF and PRC2 complex assembly.

Effective epigenetic TAZ therapy must also upregulate PRC2-repressed tumor suppressor loci (**Figure 6; Step 3**). In our patient cohort, one TAZ-resistant tumor had deletions of both *CDKN2A* and *CDKN2B* (**Figure 1A**), which can inhibit CDK4/6 from phosphorylating RB1 and are known to be de-repressed by TAZ treatment (11). Our genomic analysis of MRT and ES cell lines also showed that 4 out of 5 TAZ-resistant cell lines tested had apparent loss of *CDKN2A*, with two also having loss of *CDKN2B*, and one having *CDKN1A* loss as well (**Figure 3B**). These mutations may phenocopy *RB1* loss, suggesting that upregulation of these cell cycle inhibitors may be necessary for effective TAZ therapy. Indeed, loss of *CDKN1A* or *CDKN2A* is sufficient to confer resistance to TAZ, similar to loss of *RB1*.

Our findings establishing *CDKN2A* as a TAZ resistance mutation are particularly significant given the recent genomic studies of epithelioid sarcomas (67, 68), which found recurrent *CDKN2A* mutations with loss of p16 expression in more than one third of tumors (67). This also suggests that epigenetic dysregulation of p16 expression, for example due to repression by non-EZH2 regulators, may explain the prevalence of TAZ resistance in other *SMARCB1*-deficient tumors, such as brain ATRT and extracranial rhabdoid tumors.

In addition to maintenance of the expression of these tumor suppressor loci, transcription factors and coactivators that upregulate their expression must also be intact and expressed for tumor cells to effectively respond to TAZ (**Figure 6; Step 3**). In our patient cohort, two TAZ-resistant patient tumors had missense mutations in *ANKRD11* (**Figure 1A**), as did 3 out of 5 TAZ-resistant cell lines. *ANKRD11* is a putative tumor suppressor that exhibits loss of heterozygosity in breast cancer (69), and is recurrently mutated in other cancers (70, 71). *ANRKD11* can cooperate with p53 to upregulate *CDKN1A*, and previous reports have suggested a possible risk of cancer development in patients with a constitutional loss of *ANKRD11* (72, 73). We should note that our inability to obtain viable clones with biallelic *ANKRD11* loss, together with the presence of only missense *ANKRD11* mutations in our samples, suggests a pleiotropic function of this gene. Further work to elucidate the role of *ANKRD11* in TAZ response will require the introduction of specific missense mutations detected in cell lines and tumors that can separate the cell viability functions from any effects on TAZ resistance. Interestingly, a recent report described a patient with KBG syndrome, a developmental condition caused by mutation of *ANKRD11*, who also developed a rhabdoid tumor (73). The co-occurrence of these two exceedingly rare conditions in the same patient is consistent with potential functional involvement of *ANKRD11* in rhabdoid tumor development and susceptibility to EZH2 inhibition.

We also identified *KLF4* as a putative marker of susceptibility to TAZ in patient tumors (**Figure 3G**), and found *KLF4* to be mutated in 1 out of 5 TAZ-resistant cell lines (**Figure 3B**). Although KLF4 is a transcription factor with both activating and repressing functions, its role in *SMARCB1*-deleted tumors is not known. Its expression has long been known to upregulate *CDKN1A*, causing cell cycle arrest at the G1/S checkpoint (45, 74). This suggests that KLF4 may regulate the induction of tumor suppressor genes in response to EZH2 inhibition, such as its bona fide target *CDKN1A*. This may occur through recruitment of BAF to tumor suppressor loci; KLF4 can recruit BAF to target genes to upregulate them (75). Like *ANKRD11*, *KLF4* may thus be a key activator of PRC2-repressed genes.

Finally, effective TAZ therapy also requires the function of downstream cell cycle effectors of the relevant tumor suppressor loci (**Figure 6; Step 4**). As we have demonstrated in this study, loss of *RB1* leads to the evasion of TAZ-induced cell cycle arrest, despite effective inhibition of EZH2 activity and sustained transcriptional response to TAZ. A recent genome-wide CRISPR screen also found *RB1* as a top mediator of TAZ resistance (19). This is reminiscent of the necessity of intact *RB1* for therapeutic response to clinical inhibitors of CDK4/6 (76–78). The transcriptional upregulation of hundreds of genes by TAZ in *RB1^del^* tumors, including EMT gene sets (**Figure S2C&F**), suggests that these cells are undergoing forced differentiation, even while maintaining proliferation (**Figure 2G-H**). This upregulation of mesenchymal gene sets is consistent with previous work that has shown that PRC2 inhibition, similar to *SMARCB1* re-expression, can drive *SMARCB1*-deficient tumors into a terminally differentiated, mesenchymal-like state (11, 30, 79), possibly recapitulating the normal developmental trajectory of their cells of origin (30). In this way, *RB1* loss and dysregulation of the RB1/E2F axis appear to decouple the regulation of cell fate and identity from its control of cell cycle progression. Although in our patient tumor cohort, only a single *RB1* mutation was detected, recent genomic studies found *RB1* loss to be one of the few co-occurring mutations in primary and metastatic or recurrent epithelioid sarcomas (68). Together with the mutational deletion of CDKN2A, mutation of *ANKRD11,* and epigenetic silencing of p16, this mechanism therefore represents a frequent perturbation of the G1/S checkpoint that accounts for TAZ resistance in 5 out of 16 TAZ-resistant tumor specimens studied, as well as in 4 of 5 TAZ-resistant cell lines. Further work will be needed not only to define the contribution of other mutations and their place within the model we propose here (**Figure 6**), but also on the non-cell autonomous effects of EZH2 inhibition *in vivo*, which may include increased immunogenicity stimulated by TAZ treatment and consequent immunologic tumor clearance *in vivo* (**Figure S16**).

In our search for predictive biomarkers of TAZ response, we identified increased expression of the PCP gene *PRICKLE1* to be associated with a deficient RB1/E2F axis and TAZ resistance (**Figure 3C-F**). The molecular mechanism connecting G1/S dysregulation and *PRICKLE1* expression is currently unknown, but the PCP pathway is known to be under cell cycle control (80, 81). This is mediated at least in part by PLK1, which we found to be one of the top upregulated genes in TAZ-resistant tumors (**Figure S5B**). Likewise, deletion of the *Drosophila RB1* homologue *Rbf1* results in the upregulation of several PCP genes, including the *PRICKLE1* homologue *pk* (82). This suggests that dysregulation of the RB1/E2F axis may lead to upregulation of *PRICKLE1* through dysregulation of normal cell cycle control of PCP. Further work will be needed to define this mechanism, as well as to investigate PRICKLE1 as a clinical biomarker for TAZ resistance.

Finally, these findings enabled two strategies to circumvent clinical TAZ resistance. First, since dysregulation of the RB1/E2F axis mediates escape from cell cycle arrest at the G1/S checkpoint, cell cycle kinases that function downstream of this checkpoint should remain viable therapeutic targets. This cell cycle bypass strategy is supported by previous work showing that loss of *RB1* can sensitize cancer cells to Aurora kinase inhibition through a primed spindle assembly checkpoint (83). As predicted, *RB1^del^* cells remain sensitive to CDK2, AURKA, and AURKB inhibition (**Figures 4A, S6A-B**). We found that MRT and ES cell lines resistant to TAZ, including those with *ANKRD11* and *CDKN1A/2A/2B* mutations were sensitive to barasertib. Most compellingly, we found that the combination of TAZ and barasertib exhibits improved antitumor activity *in vivo* compared to either drug alone.

This is reminiscent of the therapeutic combination mechanism proposed by Sorger and Palmer (84), in which inter-patient and inter-tumor variability in response to individual drugs rather than their pharmacological interactions lead to apparent combined effects. We observed substantial benefit of TAZ and barasertib combination within individual PDX models. This suggests that the improved efficacy of this combination may also result from intra-tumor heterogeneity and tumor evolution *in vivo*, as proposed for combination therapy more than fifty years ago (85, 86). Dual targeting of two different parts of the cell cycle can prevent tumors from evading cell cycle arrest through the presence or acquisition of mutations in cell cycle control genes. For example, cells with a defective G1/S checkpoint that continue to transit through the cell cycle remain sensitive to AURKB inhibition. Further work will be needed to elucidate whether barasertib causes postmitotic endoreduplication due to its disruption of the spindle assembly and/or direct inhibition of the mitotic checkpoint (87, 88).

Our findings also advance a synthetic lethal strategy for rational epigenetic TAZ combination therapy due to TAZ-induced expression of *PGBD5*, the putative developmental mutator in rhabdoid and other young-onset solid tumors, through its induction of DNA damage and requirement for ongoing DNA damage repair signaling (57). We found that the DNA damage repair ATR kinase inhibitor elimusertib not only can overcome RB1/E2F axis-mediated resistance, but in combination with TAZ, can also exert synergistic anti-tumor effects *in vitro* and *in vivo*. Indeed, PGBD5 is required for the induction of DNA damage by the combination of TAZ and elimusertib, substantiating the specific nature of the synthetic lethal dependency and therapeutic vulnerability created by TAZ-mediated induction of PGBD5 and its DNA nuclease activity.

It is possible that the enhanced sensitivity to ATR inhibition due to the increased requirements for DNA repair also depends on the intrinsic variation in DNA damage repair signaling among different tumor subtypes. This may also be due to the variation in the expression and activity of PGBD5 nuclease activity among tumors, both of which may be associated with the recently described molecular subtypes of ES and MRT tumors (56). Finally, additional synthetic lethal strategies may also be developed based on the immunologic effects of EZH2 inhibitors, particularly as combined with epigenetic and synthetic lethal therapies, both of which can promote tumor immunogenicity. In all, this study develops a paradigm for rational epigenetic combination therapy, including candidate prognostic biomarkers, all of which should be incorporated into future clinical trials for patients.

## Supporting information

Uncropped Western Blot

## Acknowledgements

This work is dedicated to Maggie Schmidt and her family, and many other patients and their advocates who inspire and support our research. We thank Richard Koche, Nicholas Socci, and Mithat Gönen for technical advice, Alejandro Gutierrez and Stuart Orkin for *EZH2* expression vectors, members of our labs for critical advice and manuscript comments, and Epizyme (now Ipsen), Bayer and AstraZeneca for supplying tazemetostat, elimusertib, and barasertib, respectively. This work was supported by the MSK Integrated Genomics Operation Core, Anti-Tumor Assessment Core, Bioinformatics Core, Molecular Diagnostics Service and the Department of Pathology, the Marie-Josée and Henry R. Kravis Center for Molecular Oncology, NIH R01 CA214812, P30 CA08748, T32 GM007739, Burroughs Wellcome Fund, Rita Allen Foundation, Pershing Square Sohn Cancer Research Alliance and the G. Harold and Leila Y. Mathers Foundation, Cycle for Survival, MSK Sarcoma Center, the Starr Cancer Consortium, and Maggie’s Mission. Yaniv Kazansky was supported by a Medical Scientist Training Program grant from the National Institute of General Medical Sciences of the National Institutes of Health under award number T32 GM007739 to the Weill Cornell/Rockefeller/Sloan Kettering Tri-Institutional MD-PhD Program. AK is a Scholar of the Leukemia & Lymphoma Society.

## Methods

### *EZH2* mutant plasmids

The *EZH2^Y666N^* mutation detected in the clinical trial patient refers to amino acid numbering in isoform 2 of the protein. For consistency of nomenclature, all engineered mutations use numbering referring to isoform 2 (Uniprot ID: Q15190-2), although isoform 1 was expressed in cells for this study. Plasmids containing wild-type EZH2 (*EZH2^WT^*) and catalytically inactive triple mutant (F672I, H694A, R732K, referred to as *EZH2^CatMut^*) plasmids were kindly provided by Alejandro Gutierrez in the doxycycline-inducible pINDUCER20 vector (89). The plasmids contain human *EZH2* tagged N-terminally with a FLAG-Avi tag.

The Y666N mutation was engineered in both the *EZH2^WT^*and *EZH2^CatMut^* plasmids to yield *EZH2^Y666N^* and what we termed *EZH2^QuadMut^*. The mutation was introduced by site-directed mutagenesis using the QuikChange Lightning Kit (Agilent) per manufacturer’s instructions (mutagenesis primers: 5’-GCAAAGTGTACGACAAGAACATGTGCAGCTTTCTG-3’ and 5’-CAGAAAGCTGCACATGTTCTTGTCGTACACTTTGC-3’) to engineer a TAC to AAC codon change. After mutagenesis, PCR products were transformed into Stbl3 *E. coli* cells and expanded at 30°C. Correct plasmid sequences were confirmed by Sanger sequencing (Eton). See oligonucleotide table below for all sequencing primers used.

### Generating CRISPR knockout cells

G401 cells with *RB1* mutations were engineered by Synthego. Briefly, a single guide RNA targeting exon 2 of *RB1* was used (see oligonucleotide table below for gRNA and primer sequences). Ribonucleoprotein containing spCas9 and sgRNA was transfected into G401 cells by electroporation. The target site was then PCR-amplified and Sanger sequenced to ensure homozygous indels. The cells were then single-cell cloned and re-verified by Sanger sequencing. Loss of the RB1 protein was confirmed by Western blot.

G401 cells with *CDKN2A* and *CDKN1A* KO were generated in-house, using crRNAs against exon 1 of *CDKN2A* and exon 2 of *CDKN1A*. Ribonucleoproteins were prepared using crRNA, ATTO550-labeled tracrRNA, and Alt-R S.p. Cas9 (IDT). Cells were transfected by electroporation using the Neon transfection system (Invitrogen) with parameters set at 1300 V, 20 ms, 1 pulse Transfected cells were selected by flow sorting and single-cell cloned. Homozygous indels were confirmed by Sanger sequencing (see table below for gRNA and primers sequences).

### Generating shPGBD5 cells

For shRNA cells, pLKO.1 shRNA vectors targeting *PGBD5* (TRCN0000138412, TRCN0000135121) and control shGFP were obtained from the RNAi Consortium (Broad Institute). G401 cells were transduced at an MOI ∼1.5 and selected with puromycin at 2 µg/mL for 72 hours. Knockdown was confirmed by quantitative RT-PCR (see Supplementary Methods) using primers specified in the oligonucleotide table below.

#### Oligonucleotides

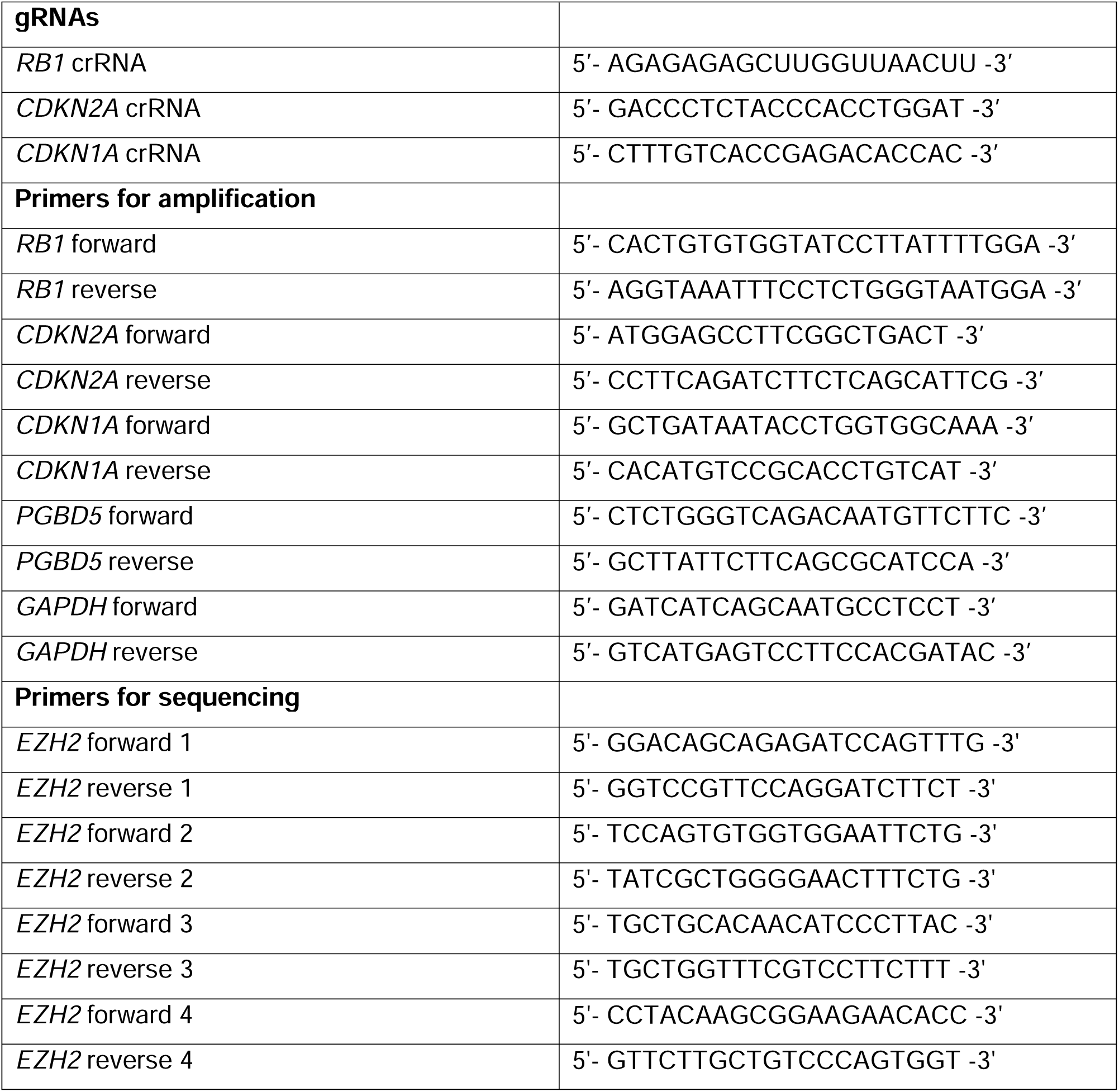
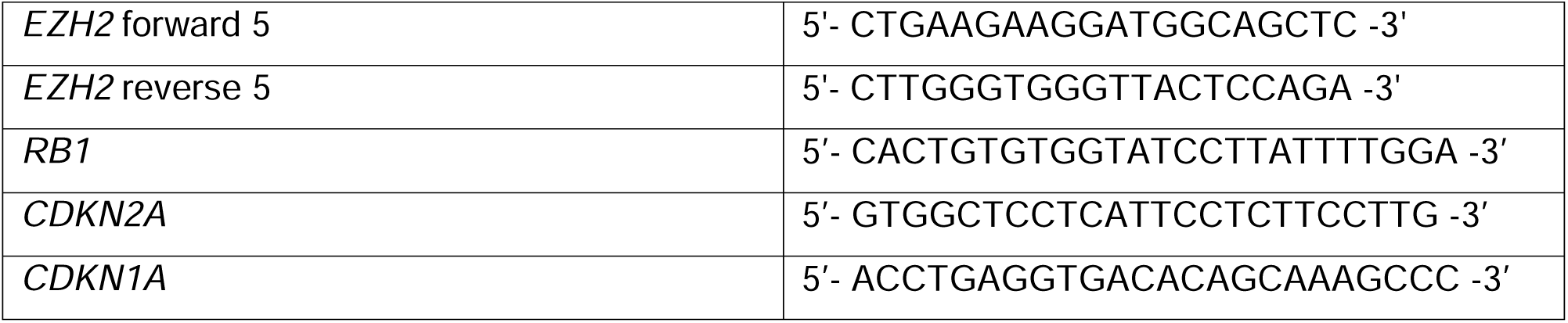

### Cell cycle analysis

G401 cells were plated on Day 0 and treated for 10 days with 1 µm tazemetostat or equivalent volume of DMSO, with drug and media replaced on Days 4 and 7. On Day 11, cells were pulsed with EdU for 1 hour. Cells were then harvested and processed for flow cytometry using the manufacturer’s protocol (Click-iT, Invitrogen). Briefly, cells were washed with PBS with 1% BSA, permeabilized, and incubated with AlexaFluor647 for 30 minutes. DNA content was measured using propidium iodide (0.05 µg/µL). Cells were analyzed on a CytoFLEX LX (Beckman Coulter).

### Patient tumor samples

Patient tumor and matched normal blood samples were obtained from patients at Memorial Sloan Kettering Cancer Center (MSKCC) enrolled in the TAZ clinical trial (15). All patients provided informed consent for this study under the Institutional Review Board approved research protocol 12-245. Patient tumors were classified into “Response” or “Progression” groups based on RECIST 1.1 criteria (90). “Response” included tumors exhibiting a complete response, partial response, or stable disease. All other tumors were classified under the “Progression” group. The complete list of tumor samples used and corresponding clinical data may be found in Supplementary Table 1. This cohort includes both tumor samples that underwent targeted sequencing with MSK-IMPACT (20) as part of their clinical care at MSKCC as well as archived tumors that were analyzed for this study. For genomic analysis, DNA was extracted from either flash frozen tumor samples or formalin-fixed, paraffin-embedded (FFPE) blocks or slides and samples were processed using the IMPACT468 or IMPACT505 panels depending on the time of their sequencing (20). The detected mutations and copy number alterations were obtained from cBioPortal (91, 92) and can be found in Supplementary Table 2. Oncoprints were generated using Oncoprinter (cBioPortal).

For transcriptomic analysis, archived frozen tumor samples were weighed and up to 20-30mg were homogenized in RLT buffer, followed by extraction using the AllPrep DNA/RNA Mini Kit (QIAGEN catalog 80204) according to the manufacturer’s instructions. RNA was eluted in 13 µL nuclease-free water. After RiboGreen quantification and quality control by Agilent BioAnalyzer, 1 µg of total RNA with DV200 percentages varying from 78% to 100% underwent ribosomal depletion and library preparation using the TruSeq Stranded Total RNA LT Kit (Illumina catalog RS-122-1202) according to instructions provided by the manufacturer with 8 cycles of PCR. Samples were barcoded and sequenced using NovaSeq 6000 in a PE150 mode, with the NovaSeq 6000 S4 Reagent Kit (Illumina). On average, 84 million paired reads were generated per sample and 70% of the data mapped to mRNA.

### Targeted sequencing of cell lines

To assess for the presence of somatic mutations in MRT and ES cell lines, DNA was extracted using the PureLink Genomic DNA Minikit (Invitrogen) and processed using the IMPACT505 panel as above. Due to the lack of matched normal tissue for cell lines, copy number alterations were detected using a custom algorithm using circular binary segmentation (93) implemented by the MSK Bioinformatics core. Code is available on github at: https://github.com/kentsisresearchgroup/seqCNA_tazemetostat_resistance.

Additional methods may be found in the Supplementary Methods and Figures file.

## Data Availability

RNA-seq data from G401 cells can be found at the Gene Expression Omnibus (GEO) repository, accession number GSE213845. RNA-seq data from patient tumor samples has been deposited to the Database of Genotypes and Phenotypes (dbGaP), accession number phs003188.v1.p1. Additional supplementary data files and computational analysis scripts are available at Zenodo (https://doi.org/10.5281/zenodo.7595037).

## Supplementary methods

### Cell culture

All cell lines were obtained from the American Type Culture Collection if not otherwise specified. ES1 and ES2 cells were generated and kindly provided by Nadia Zaffaroni. EPI544 cells were obtained from the MD Andersen Cancer Center Cytogenetics and Cell Authentication Core. Rhabdoid tumor cell lines KP-MRT-NS, KP-MRT-RY, and MP-MRT-AN were kindly provided by Yasumichi Kuwahara and Hajime Hosoi. The identity of all cell lines was verified by STR analysis. Absence of *Mycoplasma* contamination was determined using the MycoAlert kit according to manufacturer’s instructions (Lonza). Cell lines were cultured in 5% CO_2_ in a humidified atmosphere in 37°C. All media were obtained from Corning and supplemented with 10% fetal bovine serum (FBS), 1% L-glutamine, and 100 U/mL penicillin and 100 µg/mL streptomycin (Gibco). RPE, G401, A204, ES1, ES2, and VAESBJ cells were cultured in Dulbecco’s Modified Eagle Medium (DMEM). TTC642, TM8716, MP-MRT-AN, KP-MRT-NS, and KP-MRT-RY cells were cultured in Roswell Park Memorial Institute (RPMI) medium. EPI544 cells were cultured in DMEM/F12 medium.

### Western Blotting

To assess protein expression by Western immunoblotting, pellets of 1 million cells were prepared and washed once in cold PBS. Cells were resuspended in 100-130 µL of RIPA lysis buffer (50 mM Tris-HCl, pH 8.0, 150 mM NaCl, 1.0% NP-40, 0.5% sodium deoxycholate, 0.1% sodium dodecyl sulfate) and incubated on ice for 10 minutes. Cell suspensions were then disrupted using a Covaris S220 adaptive focused sonicator for 5 minutes (peak incident power: 35W, duty factor: 10%, 200 cycles/burst) at 4 °C. Lysates were cleared by centrifugation at 18,000 g for 15 min at 4 °C. Protein concentration was assayed using the DC Protein Assay (Bio-Rad) and 15-35 µg whole cell extract was used per sample. Samples were boiled at 95 °C in Laemmli buffer (Bio-Rad) with 40 mM DTT and resolved using sodium dodecyl sulfate-polyacrylamide gel electrophoresis. Proteins were transferred to Immobilon FL PVDF membranes (Millipore), and membranes were blocked using Intercept Blocking buffer (Li-Cor). Primary antibodies used were: anti-EZH2 (Cell Signaling Technology, 5246) at 1:1,000, anti-RB1 (Cell Signaling Technology, 9309) at 1:250, anti-H3K27me3 (Cell Signaling Technology, 9733) at 1:500, anti-p16 (Abcam, ab108349) at 1:500, anti-CCNA2 (Santa Cruz, sc-271682) at 1:100, anti-PRICKLE1 (Santa Cruz, sc-393034) at 1:100, anti-SMARCB1 (BD Biosciences, 612110) at 1:500, anti-RPA32 pT21 (abcam, ab109394) at 1:2,000, anti-RPA32 pS4/pS8 (ThermoFisher, A300-245A) at 1:2,000, anti-pCHK1 S296 (Cell Signaling Technology, 90178) at 1:250, anti-MYCN (Cell Signaling Technology, 9405) at 1:250, anti-MMP2 (Cell Signaling Technology, 40994S) at 1:250, anti-cleaved Caspase 3 (Cell Signaling Technology, 9661S) at 1:500, anti-Actin (Cell Signaling Technology, 4970 and 3700) at 1:5,000. Blotted membranes were visualized using goat secondary antibodies conjugated to IRDye 680RD or IRDye 800CW (Li-Cor, 926-68071 and 926-32210) at 1:15,000 and the Odyssey CLx fluorescence scanner, according to manufacturer’s instructions (Li-Cor). Image analysis was done using the Li-Cor Image Studio software (version 4).

### Lentivirus production

Lentivirus production was carried out as described previously (1). Briefly, HEK293T cells were transfected using TransIT-LT1 using a 2:1:1 ratio of the lentiviral vector and psPAX2 and pMD2.G packaging plasmids, according to manufacturer’s instructions (Mirus). Viral supernatant was collected at 48 and 72 hours post-transfection, pooled, filtered and stored in aliquots at -80 °C. G401 cells were transduced at a multiplicity of infection (MOI) of 0.3. Transduced cells were selected for 7 days with G418 sulfate (ThermoFisher) at 1 mg/mL. Single-cell clones were then isolated and expanded. Inducible EZH2 expression was confirmed by Western blotting against EZH2.

### Cell Viability Testing

Drugs used for *in vitro* treatment were supplied by Selleckchem (TAZ; S7128, Elimusertib; S9864, abemaciclib; LY2835219, palbociclib; S1116, seleciclib; S1153, alisertib; S1133, barasertib; S1147, camptothecin, S1288).

The effects of *RB1* loss or *EZH2* mutation on TAZ susceptibility was assessed over 14 days. Cells were plated in 96-well microplates at equal densities and treated with 10 µM TAZ or equivalent volume of DMSO on Day 0. Drug and media were replaced on Days 4, 7, and 11. CellTiter-Glo assays were performed on Day 14, with luminescence readings taken using an automated fluorescence plate reader (Tecan). CellTiter-Glo reagent was freshly reconstituted on the day of measurement and added in a 1:1 proportion to cell media. A similar protocol was used for all other cell viability experiments, with treatment times indicated in the relevant figure legends. Cell line doubling time was determined by measuring cell viability every 24 hours over the course of 4 days, and fitting the cell viability to a two-parameter exponential curve. For combination treatment with TAZ and elimusertib, we used a two-dimensional dose matrix design, treating the cells for 9 days. After the addition of cells, drugs were added using a pin tool (stainless steel pins with 50 nL slots, V&P Scientific) mounted onto a liquid handling robot (CyBio Well vario, Analytik Jena). For analysis of synergy, we used the synergyFinder package (2). Outliers due to pinning errors were excluded after manual examination.

### Cell line RNA-sequencing

G401 cells with or without *RB1* loss were plated and treated with 10 µM TAZ or equivalent volume of DMSO on Day 0. Drug and media were replaced on Days 4 and 8. Cells were harvested on Day 11 and RNA was isolated using RNeasy Mini kit, according to manufacturer’s instructions (Qiagen). After RiboGreen quantification and quality control by Agilent BioAnalyzer, 149-500ng of total RNA underwent Poly(A) selection and TruSeq library preparation according to instructions provided by Illumina (TruSeq Stranded mRNA LT Kit, catalog RS-122-2102), with 8 cycles of PCR. Samples were barcoded and sequenced using a HiSeq 4000 instrument using 50bp/50bp paired end mode, using the HiSeq 3000/4000 SBS Kit (Illumina). An average of 42 million paired reads was generated per sample. Ribosomal reads represented less than 0.03% of the total reads generated and proportion of mRNA bases averaged 74%.

### Analysis of RNA-seq data

For RNA-seq analysis of G401 cell lines, read adaptors were trimmed and quality filtered using ‘trim_galore’ (v0.4.4_dev) and mapped to GRCh38/hg19 reference genome using STAR v2.6.0a with default parameters (3). Read counts tables were generated using HTSeq (4). Normalization was performed using DESeq2 using the default parameters (5). Differential gene expression analysis of transposable elements (TEs) was performed using the TEtranscripts (6) package with default parameters, with GTF files for TE annotations for hg19 generated by the Hammell lab using UCSC Repeat Masker.

For RNA-seq analysis of patient tumor samples, read adaptors were trimmed and quality filtered using ‘trim_galore’ and mapped to GRCh38/hg19 reference genome using STAR v2.7.9 with default parameters (3). Read count tables were generated using HTSeq v0.11.3 (4). Bam files were sorted by name using ‘samtools’ and alignment quality was assessed using ‘qualimap’ v2.2.2. Normalization was performed using DESeq2 v1.34.0 using the default parameters (5). To assess gene expression changes between TAZ-sensitive and TAZ-resistant tumors, samples in both categories were compared by two-tailed Student’s t-test using ‘rowttests’ in R v4.1.3. Genes were filtered by p<0.05 and sorted by t-statistic. Heatmaps were then generated using ‘pheatmap.’ Genome browser tracks were visualized from bam files using Integrated Genomics Viewer v2.13.1.

### Gene ontology analysis

Genes significantly up-or down-regulated in TAZ-sensitive and TAZ-resistant tumors determined by two-tailed Student’s t-test at p-value < 0.5 were searched against the Gene Ontology database (DOI: 10.5281/zenodo.5725227 Downloaded 2021-11-16).

### Microscopy

Bright field microscopy was performed using an Evos FL Auto 2 imager at 10x magnification, with cells grown on plastic dishes. Immunofluorescence for γH2AX was performed on cells plated on Millicell EZ Slide glass slides (EMD Millipore), coated for 45 minutes with bovine plasma fibronectin (Millipore Sigma). After drug treatment, cells were washed once with PBS and fixed in 4% formaldehyde for 10 minutes at room temperature. Slides were then washed three times in PBS for 5 minutes, permeabilized for 15 minutes in 0.3% Triton X-100, washed again in PBS three times, and blocked with 5% goat serum (Millipore Sigma, G9023) in PBS for 1 hour at room temperature. Slides were incubated with mouse anti-γH2A.X primary antibody (Sigma-Aldrich, 05-636) at 1:500 in blocking buffer for 1 hour, washed three times in PBS, and incubated with goat anti-mouse secondary antibody conjugated to AlexaFluor555 (Invitrogen, A-21422) at 1:1,000. Cells were then counterstained with DAPI at 1:1,000 for 10 minutes and treated with ProLong Diamond Antifade Mountant with DAPI (Invitrogen, P36962) for 48 hours. For MMP2 and cleaved caspase 3 immunofluorescence, cells were processed as above, using anti-MMP2 antibody (Cell Signaling Technology, 40994S) at 1:200 or anti-cleaved Caspase 3 antibody (Cell Signaling Technology, 9661S) at 1:300, and Phalloidin conjugated to AlexaFluor488 (ThermoFisher, A12379) at 1:400 added to the secondary antibody mix.

Images were acquired on a Zeiss LSM880 confocal microscope at 63x magnification. Images were then processed using a custom pipeline in CellProfiler (7). Per-cell integrated γH2A.X intensity was normalized against per-cell integrated DAPI intensity. All image analysis used the same pipeline settings, with the exception of the RescaleIntensity module for the AF555 channel, which used the settings 0.009-0.09 for the images in Figure 5 and 0.005-0.09 for Figure S13. Overlaid images were prepared using Fiji (8).

### Xenografts

All mouse experiments were carried out in accordance with institutionally approved animal use protocols. To generate PDXs, tumor specimens were collected under approved IRB protocol 14-091, immediately minced and mixed (50:50) with Matrigel (Corning, New York, NY) and implanted subcutaneously in the flank of 6-8 weeks-old female NOD.Cg-*Prkdc^scid^ Il2rg^tm1Wjl^/Szj* (NSG) mice (Jackson Laboratory, Bar Harbor, ME), as described previously (9). Mice were monitored daily and PDX samples were serially transplanted three times before being deemed established. PDX tumor histology was confirmed by review of H&E slides and direct comparison to the corresponding patient tumor slides. PDX identity was further confirmed by MSK-IMPACT sequencing analysis.

Therapeutic studies used female and male NSG mice obtained from the Jackson Laboratory. Xenografts were prepared as single-cell suspensions, resuspended in Matrigel, and implanted subcutaneously into the right flank of 6-10 week old mice. 100 µL of tumor cell suspension was used for each mouse. Tumors were allowed to grow until they reached a volume of 100 mm^3^, at which point they were randomized into treatment groups without blinding. Drugs were prepared using the following formulations: Tazemetostat was dissolved at 25 mg/mL in 5% DMSO, 40% PEG 300, 5% Tween 80, and 50% water. Elimusertib was dissolved at 5 mg/mL in 10% DMSO, 40% PEG 300, 5% Tween 80, and 45% water using a sonicator. Barasertib was dissolved at 2.5 mg/mL in 5% DMSO, 40% PEG 300, 5% Tween 80, and 50% water. Drugs were reconstituted daily. The following drug doses and schedules were used: TAZ was dosed at 250 mg/kg twice daily by oral gavage, 7 days per week. Barasertib was dosed at 25 mg/kg once daily by intraperitoneal injection using 3 days on and 4 days off cycle. Elimusertib was dosed at 40 mg/kg twice daily by oral gavage using 2 days on and 12 days off cycle. Caliper tumor measurements were taken twice weekly. Tumor volumes were calculated using the formula Volume = (π/6) x length x width^2^. Tumor growth analysis was performed using the Vardi *U*-test (10), as implemented in the clinfun R package using the aucVardiTest function. Tumor-free survival analysis was calculated using OriginPro (Microcal) by the Kaplan-Meier method, using the log-rank test.

### Immunohistochemistry

The immunohistochemistry detection of Ki67, Cl-caspase3, p-H2AX, p16, p21, and H3K27me3 was performed at Molecular Cytology Core Facility of Memorial Sloan Kettering Cancer Center using Discovery Ultra processor (Ventana Medical Systems). For all markers with the exception of H3K27me3, paraffin-embedded tissues were sectioned at 5 μm and baked at 58°C for 1 hr. Slides were loaded in Ultra autostainer and IHC staining was performed as follows. Samples were baked and dewaxed followed by pretreatment with epitope retrieval CC1 solution (Ventana, 950-500) for 48 min at 100°C. The primary antibodies (rabbit monoclonal Ki67 antibody (1ug/ml, Cell Signaling Technology, 9027), rabbit anti-Cleaved-caspase3 antibody (0.2ug/ml, Cell Signaling Technology, 9661), rabbit anti-p-H2AX (0.2ug/ml, Abcam, ab11174), rabbit anti p21 (0.8ug/ml, Cell Signaling Technology, 2947), and mouse anti-p16 antibody (0.2ug/ml,Santa Cruz,sc56330) were incubated for 4h at room temperature. Samples were then incubated with either OmniMap anti-rabbit HRP (Roche Diagnostics, 760-4311) secondary antibody or OmniMap anti-mouse HRP (Roche Diagnostics760-4310) for 20 minutes. Discovery ChromoMap DAB detection kit (Roche Diagnostics) was used according to the manufacturer’s instructions.

For H3K27me3 staining, slides were loaded into Leica Bond RX and dewaxed at 72°C before being pretreated with EDTA-based epitope retrieval ER2 solution (Leica, AR9640) for 20 mins. at 100°C. The Rabbit anti-H3K27me3 (0.2ug/ml, Cell Signaling Technology, 9733) were applied for 60 mins. Samples were then incubated with Leica Bond Polymer (anti-rabbit HRP) (included in Polymer Refine Detection Kit (Leica, DS9800) for another 8 min. Slides were then incubated with mixed DAB reagent (Polymer Refine Detection Kit) for 10 mins, followed by hematoxylin (Refine Detection Kit) counterstaining for 10 mins.

After staining, all sample slides were washed in water, dehydrated using an ethanol gradient (70%, 90%, 100%), washed three times in HistoClear II (National Diagnostics, HS-202), and mounted in Permount (Fisher Scientific, SP15).

**Supplementary Table S1:** List of patient tumor specimens used for RNA-seq and MSK-IMPACT analysis. *We note that two primary tumors in patients who responded to TAZ harbored deletions of *RB1* (patient 2, sample ES_02_T_02) in one tumor and *CDKN2A/B* in another tumor (patient 5, sample ES_05_T_01). However, these primary tumors were fully resected prior to the initiation of TAZ treatment and did not recur at the primary sites. In the case of patient 2, a later TAZ-responsive metastasis (ES_2_T_03) did not harbor the *RB1* loss. In the case of patient 5, a later TAZ-responsive metastasis (ES_05_T_09) did not harbor the *CDKN2A/B* loss. This suggests that these mutations were subclonal and were not present in tumors exposed to TAZ treatment. Thus, the mutations in these tumors were unlikely to have impacted their response to TAZ.

**Supplementary Table S2:** List of mutations found in all patient tumor specimens in Supplementary table 1 for which MSK-IMPACT data is available.

**Supplementary Table S3:** List of Hallmark gene sets identified by Gene Set Enrichment Analysis (GSEA) up- and down-regulated by TAZ treatment. Related to Supplementary Figure S2C.

**Supplementary Table S4:** List of mutations found in all MRT and ES cell lines used in this study as determined by targeted MSK-IMPACT sequencing. Related to Figure 3A.

**Supplementary Table S5:** List of PDX models used in this study, with clinical characteristics of the original tumor specimens, followed by a list of mutations found in all PDX models, as determined by targeted MSK-IMPACT sequencing.

**Supplementary Table S6:** List of endogenous transposable elements whose expression is up- or downregulated by TAZ treatment. Related to Supplementary Figure S16

**Figure S1:**
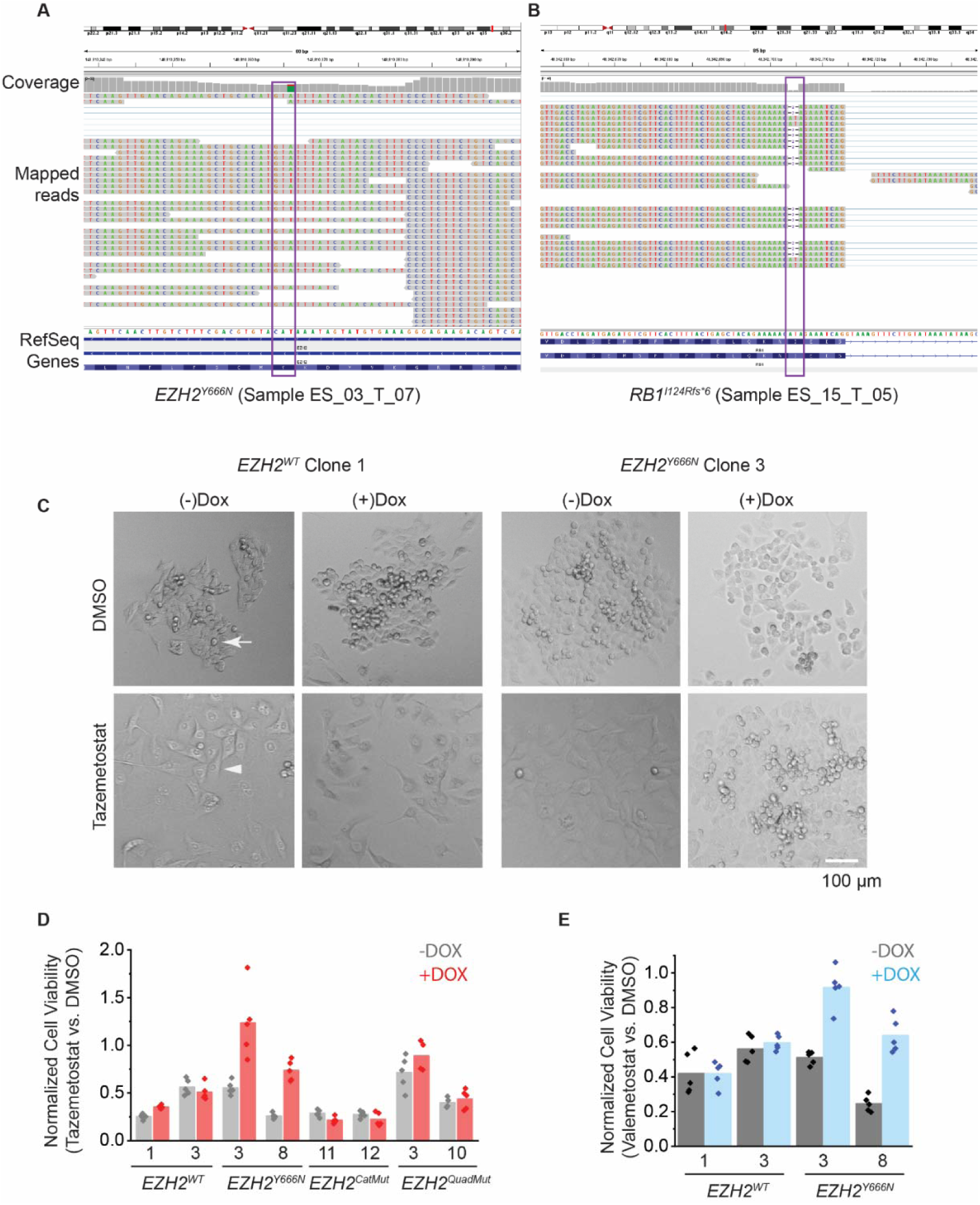
Validation of tumor resistance mutations of *EZH2*: (**A-B**) Integrated Genome Viewer (IGV) tracks of RNA-seq data for post-treatment tumor samples from patients 3 and 15. Purple boxes indicate the mutation sites. (**A**) Aligned reads and coverage plot of exon 16 of *EZH2*, showing expression of mRNA with the T (red) ➔ A (green) mutant allele. (**B**) Aligned reads and coverage plot of exon 3 of *RB1*, showing expression of mRNA with frame shift deletion at I124. (**C**) Phase-contrast microscopy of G401 single-cell clones expressing the indicated form of *EZH2*. Cells were treated with 10 µM tazemetostat or DMSO for 9 days and imaged with an Evos FL Auto 2 imager at 10X magnification. Arrow indicates a refractile, mitotic cell. Arrowhead indicates a post-treatment, morphologically altered cell. (**D**) Cell viability measured by CellTiter-Glo after 14 days of treatment with 10 µM tazemetostat or equivalent volume of DMSO. Data is the same as in Figure 1D with the addition of *EZH2^CatMut^* and *EZH2^QuadMut^* clones. *n*=5 replicates per condition. (**E**) Cell viability after treatment with 10 µM valemetostat or DMSO for 14 days. *n*=5 replicates per condition.

**Figure S2:**
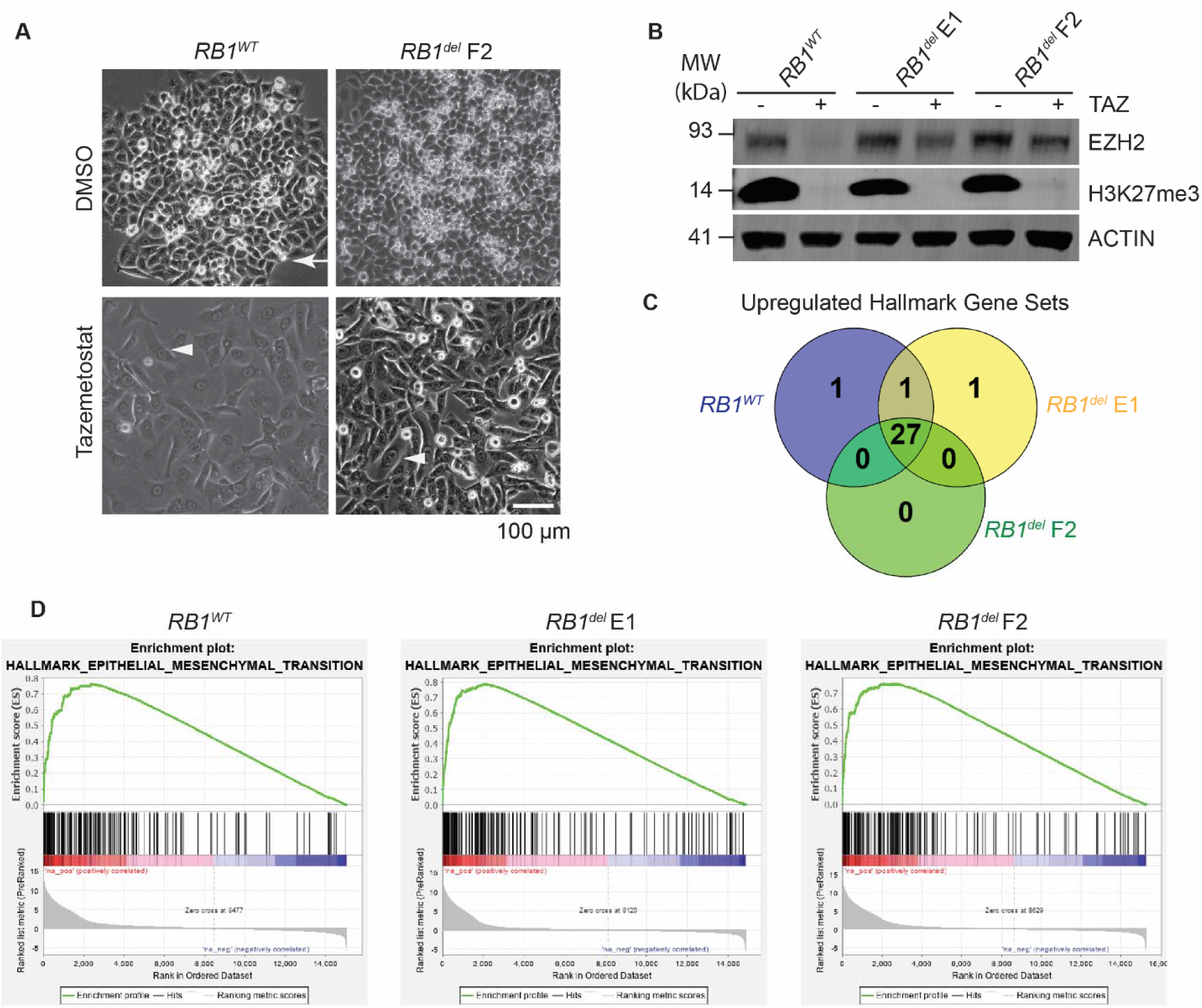
*RB1^del^*cells show morphological and transcriptional responses to TAZ: (**A**) Phase-contrast microscopy of G401 cells with or without *RB1* expression. Cells were treated with 10 µM tazemetostat or DMSO for 9 days and imaged with an Evos FL Auto 2 imager at 10X magnification. Arrow indicates a refractile, mitotic cell. Arrowhead indicates a post-treatment, morphologically altered cell. (**B**) Western blot of the indicated G401 cell clones treated with 10 µM TAZ vs. equivalent volume of DMSO for 11 days. Bulk H3K27me3 levels are reduced in all three clones despite persistent EZH2 expression in *RB1^del^* clones. (**C**) Comparison of all Hallmark gene sets upregulated in G401 cells upon TAZ treatment with significance at FDR < 25%. (**D**) GSEA plots showing the Hallmark_Epithelial_Mesenchymal_Transition gene set for the indicated TAZ-treated G401 cells compared to DMSO.

**Figure S3:**
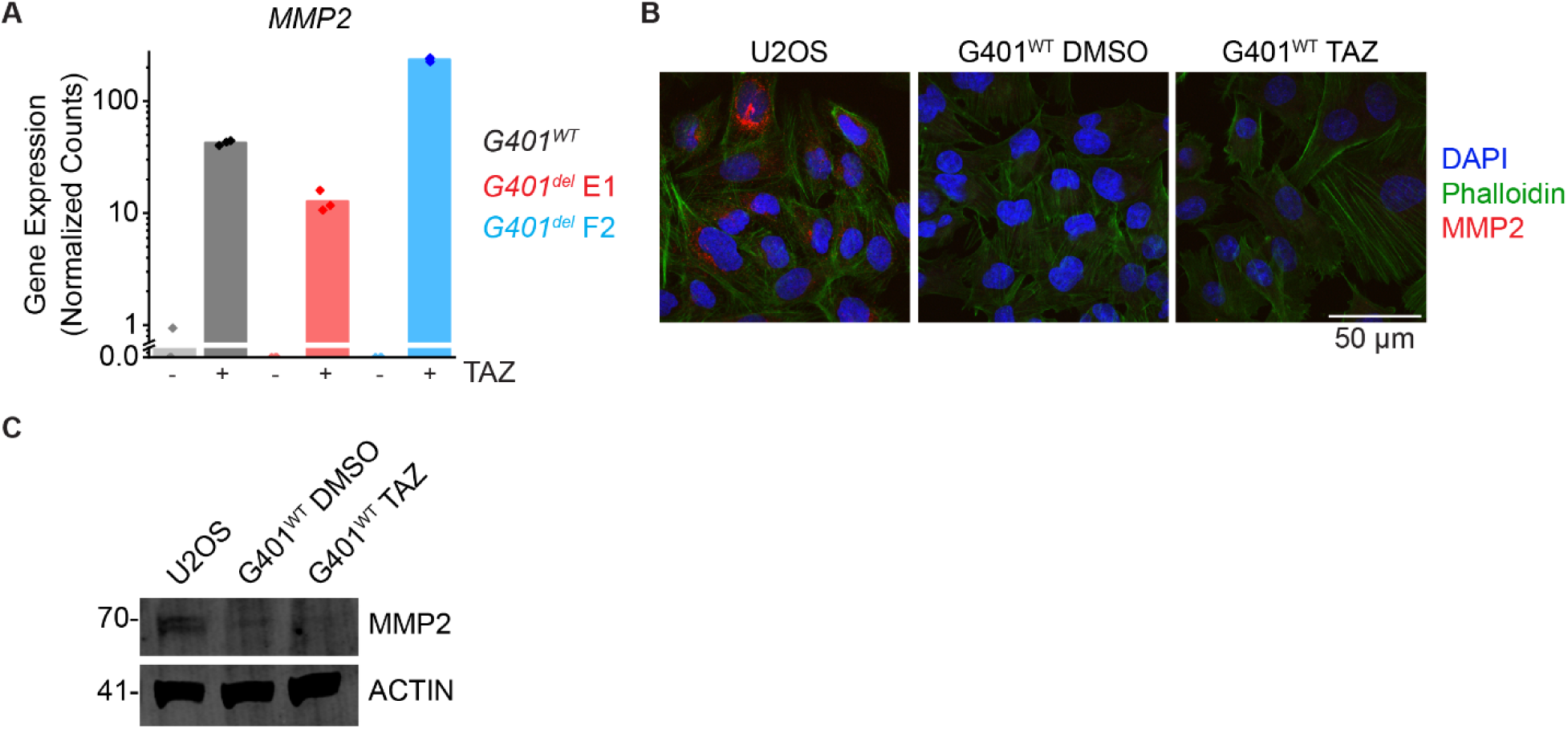
TAZ-treated *RB1^del^* cells show evidence of differentiation at the transcript, but not protein level. (**A**) RNA-seq data from cells in Figure 2A-F, showing normalized read counts for the *MMP2* gene. (**B**) Indicated cells treated with 1 µM TAZ for 11 days, stained with MMP2 antibody. U2OS cells are shown as a positive control. (**C**) MMP2 western blot on cells treated as in (B).

**Figure S4:**
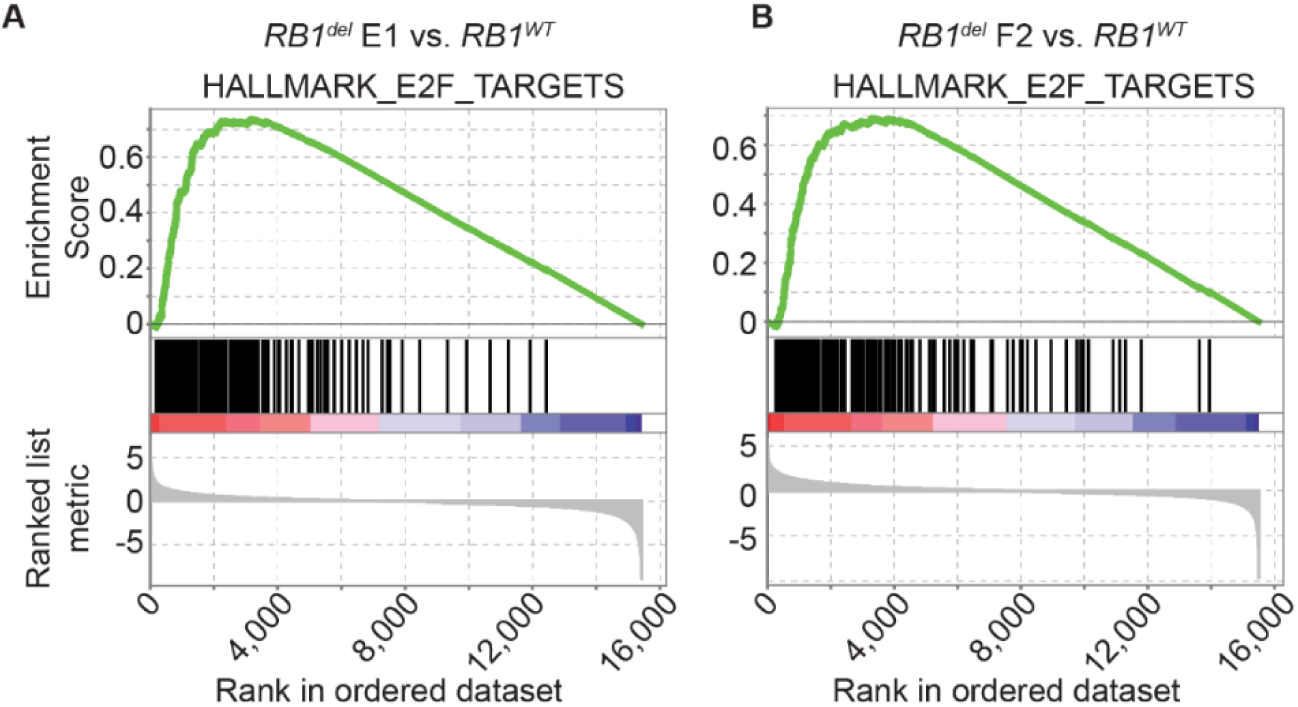
*RB1^del^* cells show increased expression of E2F targets. (**A-B**) GSEA plots showing the Hallmark_E2F_Targets gene set comparing TAZ-treated *RB1^del^* G401 cells with *RB1^WT^*cells.

**Figure S5:**
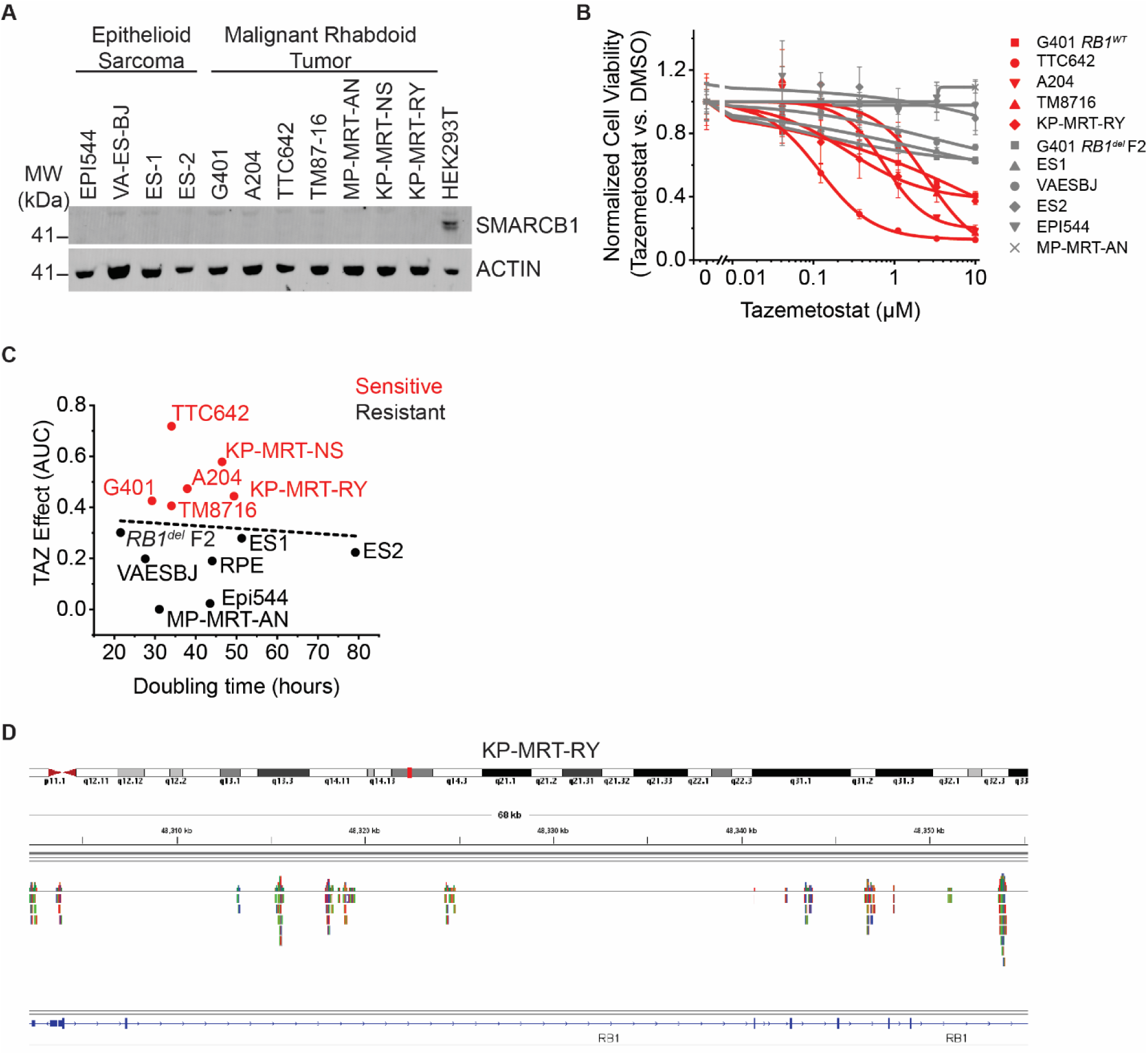
Characterization of MRT and ES cell lines: (**A**) Confirmed loss of SMARCB1 expression in all MRT and ES cell lines used. (**B**) Dose-response curves of a panel of MRT and ES cell lines treated with TAZ for 13 days. Curves in red correspond to TAZ-responsive cell lines, grey to TAZ-resistant cell lines. *n=3* replicates per condition. (**C**) Plot of area under the curve (AUC) of the dose-response curves from Figure 3A plotted against doubling time. AUC was integrated from 40 nM to 50 µM (**D**) IGV track of reads from MSK-IMPACT sequencing of KP-MRT-RY cells, focusing on the *RB1* gene, shows reads spanning segments across the full gene and indicates the presence of at least one allele.

**Figure S6:**
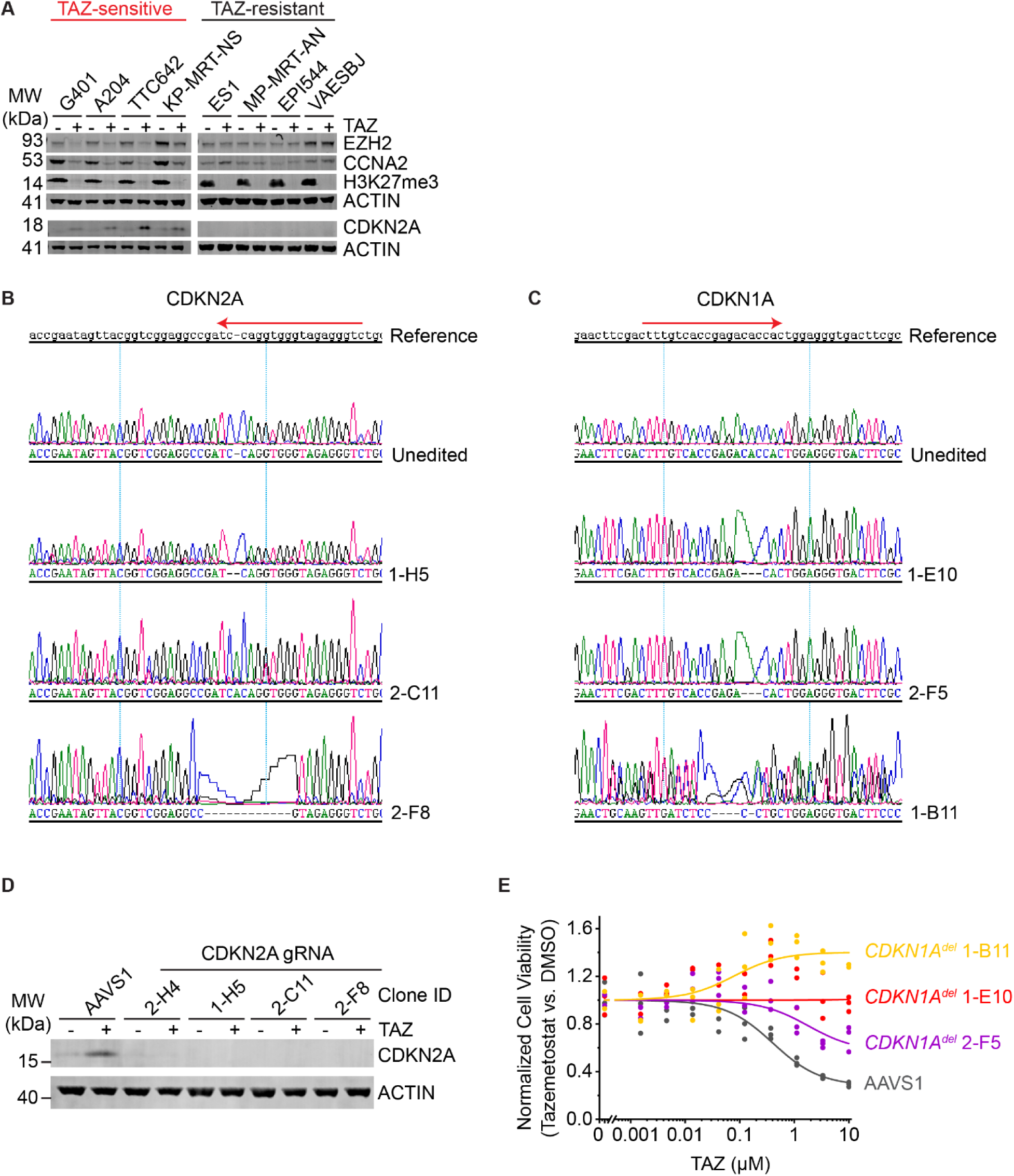
Generating and testing *CDKN1A^del^* and *CDKN2A^del^* G401 cells. (**A**) Indicated cell lines were treated with 10 µM TAZ or DMSO for 11 days. (**B-C**) Sanger traces of the indicated G401 clones for *CDKN2A^del^*(**B**) and *CDKN1A^del^* (**C**) showing homozygous editing. Red arrows indicate crRNA sequences. (**D**) Indicated G401 *CDKN2A^del^* cell clones treated with 10 µM TAZ or DMSO for 7 days, confirming loss of CDKN2A induction. (**E**) Dose response curves of the indicated cell lines treated with TAZ for 11 days. *n=3* replicates per condition.

**Figure S7:**
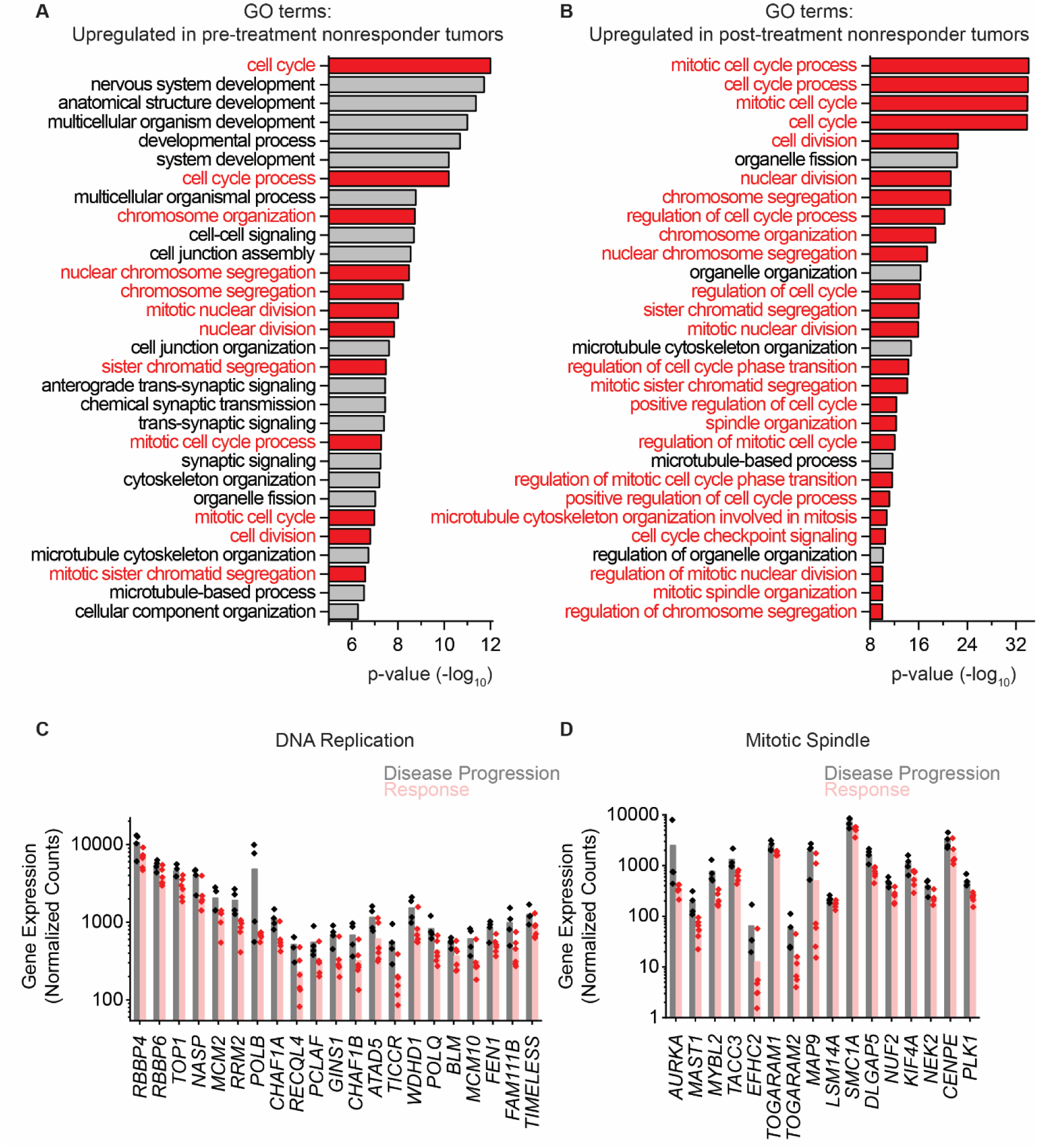
TAZ-resistant patient tumors show upregulation of cell cycle genes: Top 30 GO terms, sorted by p-value, enriched in pre-treatment (**A**) and post-treatment (**B**) patient tumor specimens that did not respond to TAZ, compared to those that did. (**C-D**) DESeq2-normalized read counts of genes from the indicated GO terms comparing pre-treatment TAZ-responding tumors to pre-treatment non-responding tumors.

**Figure S8:**
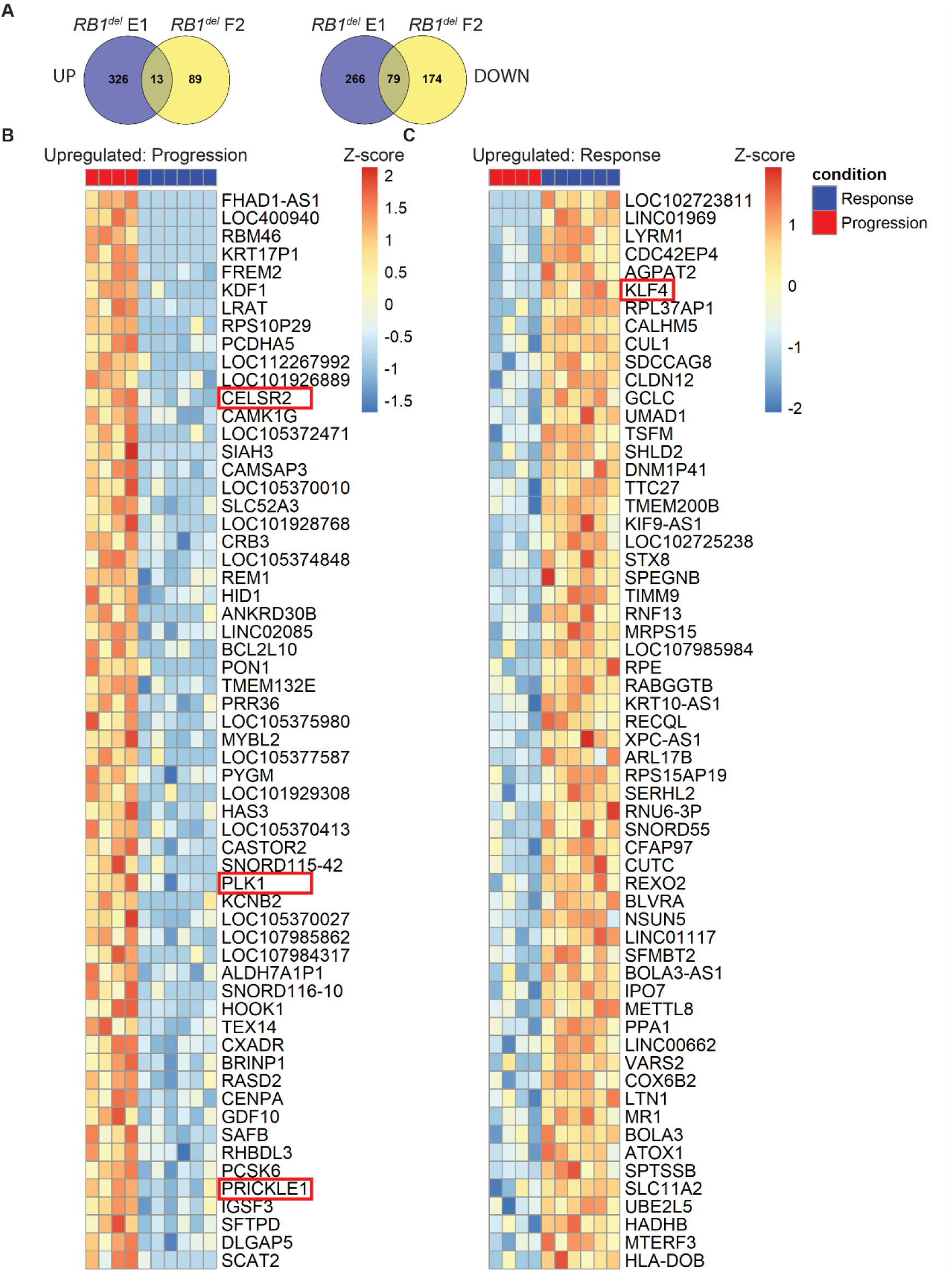
Transcriptomic analysis of patient tumors nominates putative biomarkers of TAZ sensitivity and resistance: (**A**) Venn diagrams of genes up- or down-regulated by *RB1* knockout in the indicated clone. Same data as in Figure 2. (**B-C**) Heatmaps showing the top 60 upregulated (**B**) and downregulated (**C**) genes in patient tumors collected prior to TAZ-treatment. Each column indicates a separate tumor. Heatmaps are sorted by t-statistic calculated using a two-tailed Student’s t-test, *p*<0.05. Red boxes indicate planar cell polarity genes *CELSR2*, *PLK1*, and *PRICKLE1* and *CDKN1A* regulator *KLF4*.

**Figure S9:**
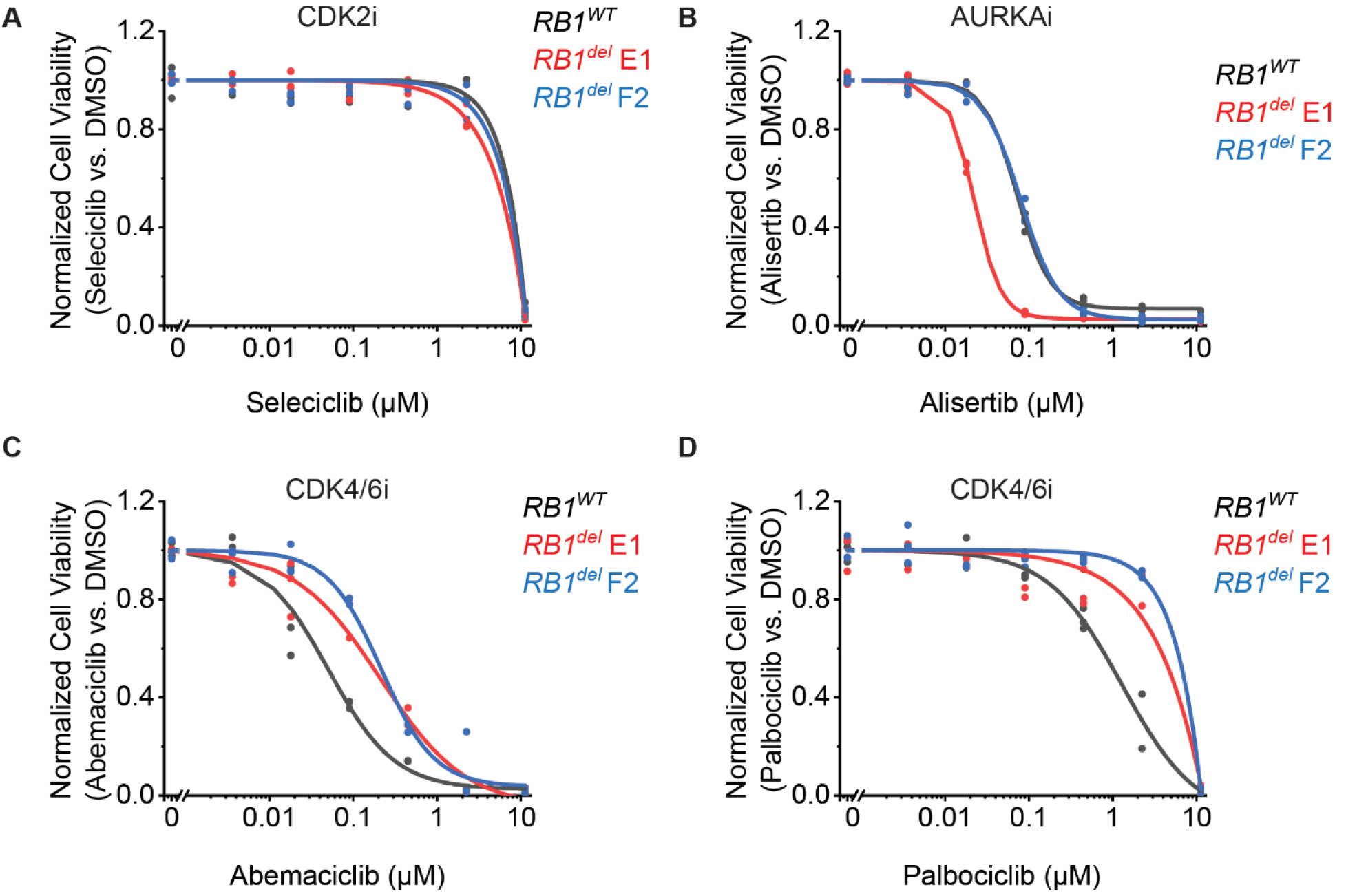
Downstream cell cycle inhibitors overcome resistance to TAZ: G401 cells treated with seleciclib (**A**), alisertib (**B**), abemaciclib (**C**), or palbociclib (**D**) 9 days. *n*=3 replicates per condition.

**Figure S10:**
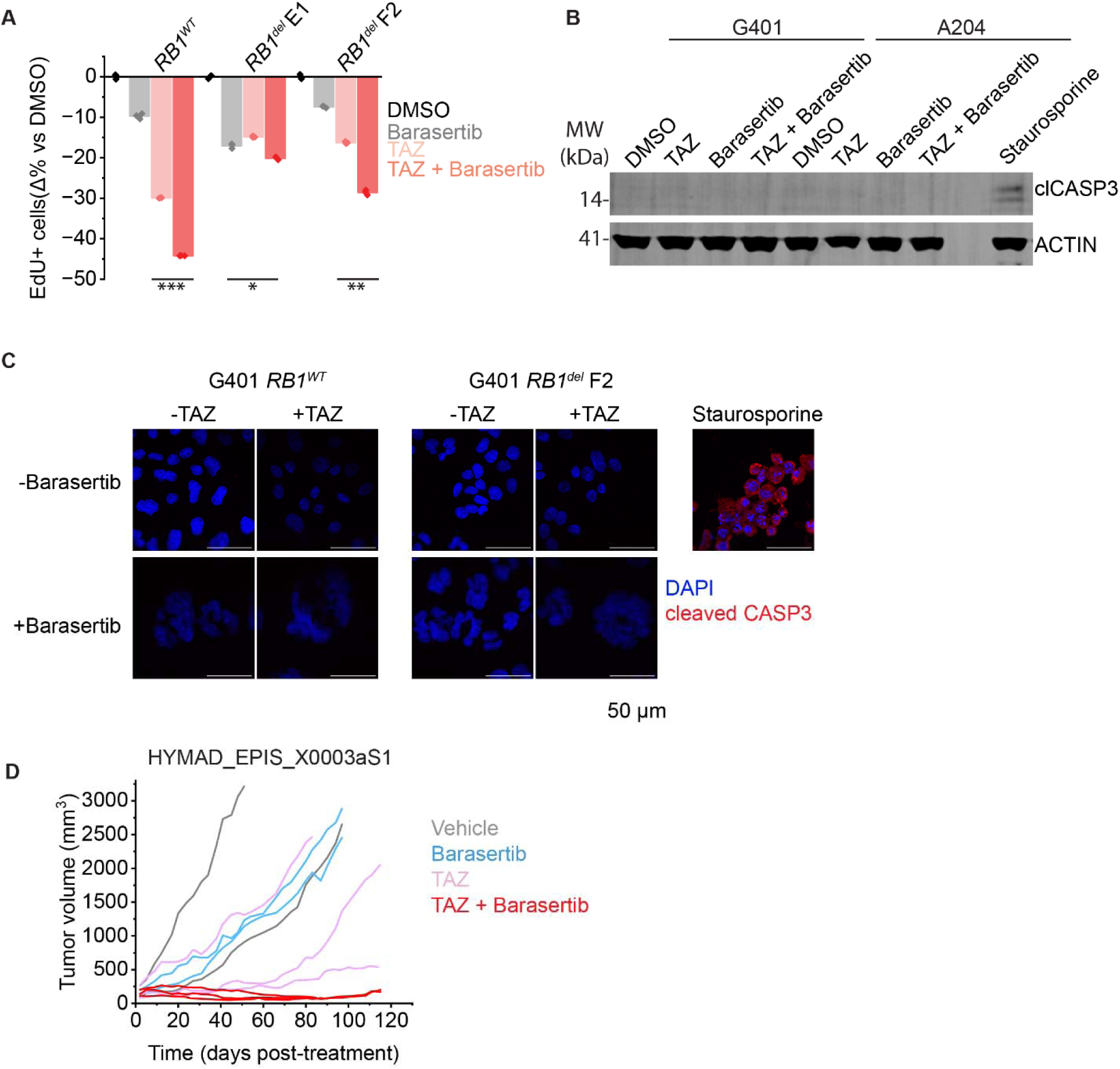
TAZ + barasertib increases cell cycle arrest without inducing apoptosis. (**A**) EdU incorporation measured by flow cytometry for the indicated cell lines and treatments. Y-axis shows percent change vs. DMSO treatment for each cell line. TAZ dose was 1 µM for 11 days, with barasertib added at 12 nM or equivalent volume of DMSO added for the last 3 days. \**p*=1.4E-3 , \*\**p*=5.3E-6 , \*\*\**p*=2.2E-9 . (**B-C**) Western blot (**B**) and immunofluorescence (**C**) of G401 and A204 cells treated as indicated, with drug doses and times as in (**A**). Staurosporine treatment for 6 h at 0.5 µM was used as a positive control. (**D**) Tumor growth curves for the subset of mouse tumors in Figure 4C from the HYMAD_EPIS_X0003aS1 PDX model. *n* = 2 mice for vehicle and barasertib-treated groups, *n* = 3 mice for TAZ and TAZ + barasertib-treated groups.

**Figure S11:**
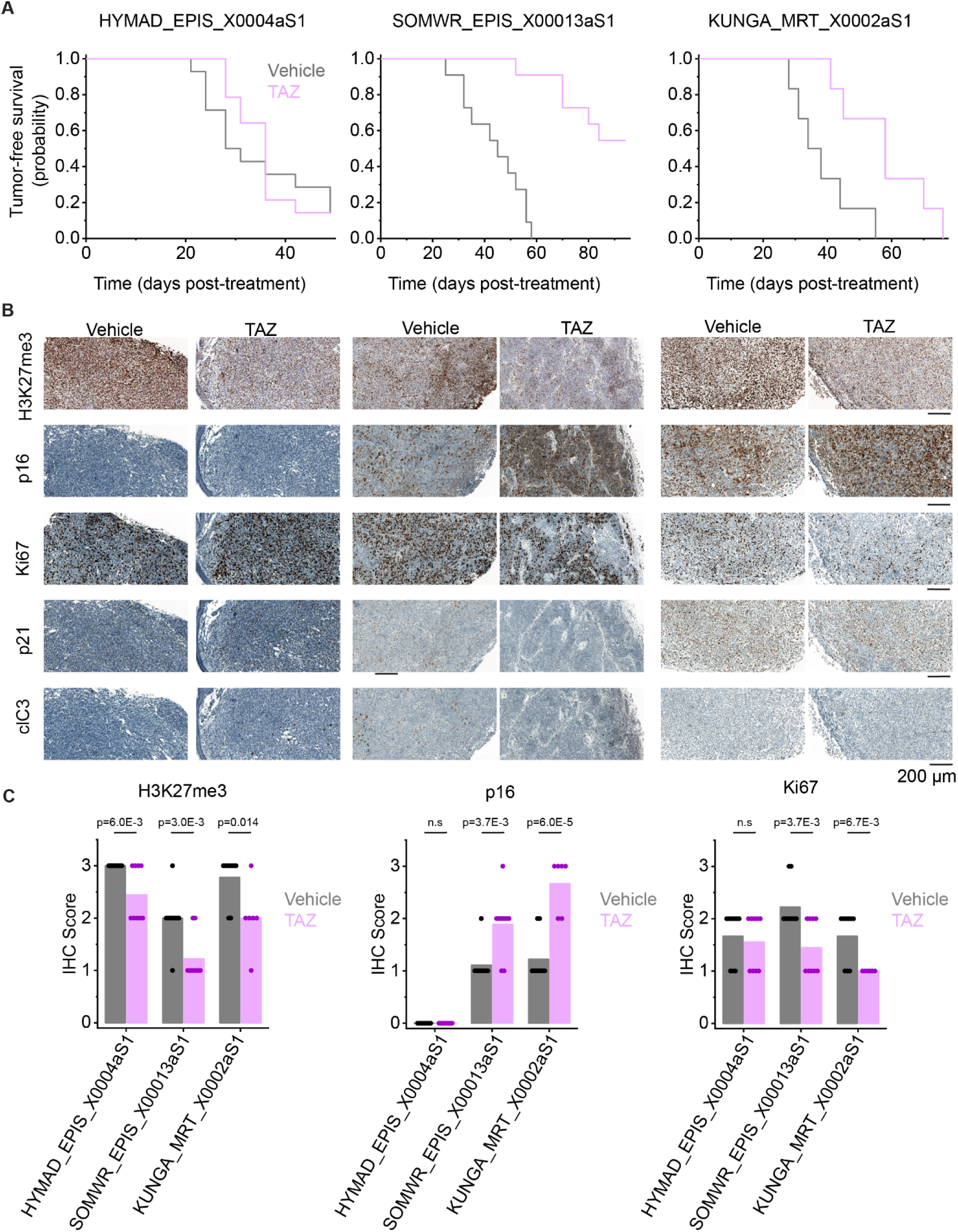
p16 induction correlates with TAZ response *in vivo*. (**A**) Kaplan-Meier curves showing tumor-free survival (defined as tumor volume ≤ 1,000 mm^3^) for the indicated PDXs. HYMAD_EPIS_X0004aS1; mean survival is 34 days (95% CI: 29-40 days) for vehicle and 36 days (95% CI: 32-39 days) for TAZ. Log-rank test *p* = 0.63. SOMWR_EPIS_X00013aS1; mean survival is 44 days (95% CI: 37-51 days) for vehicle and 84 days (95% CI: 76-92 days) for TAZ. Log-rank *p*= 7.1E-6. KUNGA_MRT_X0002aS1; mean survival is 38 days (95% CI: 30-46 days) for vehicle and 58 days (95% CI: 47-69 days) for TAZ. Log-rank *p* = 8.0E-3. (**B**) Representative IHC images of PDX tissue after treatment with vehicle vs. TAZ. (**C**) Quantification of IHC images. 3 fields for 3 tumors were quantified for each condition.

**Figure S12:**
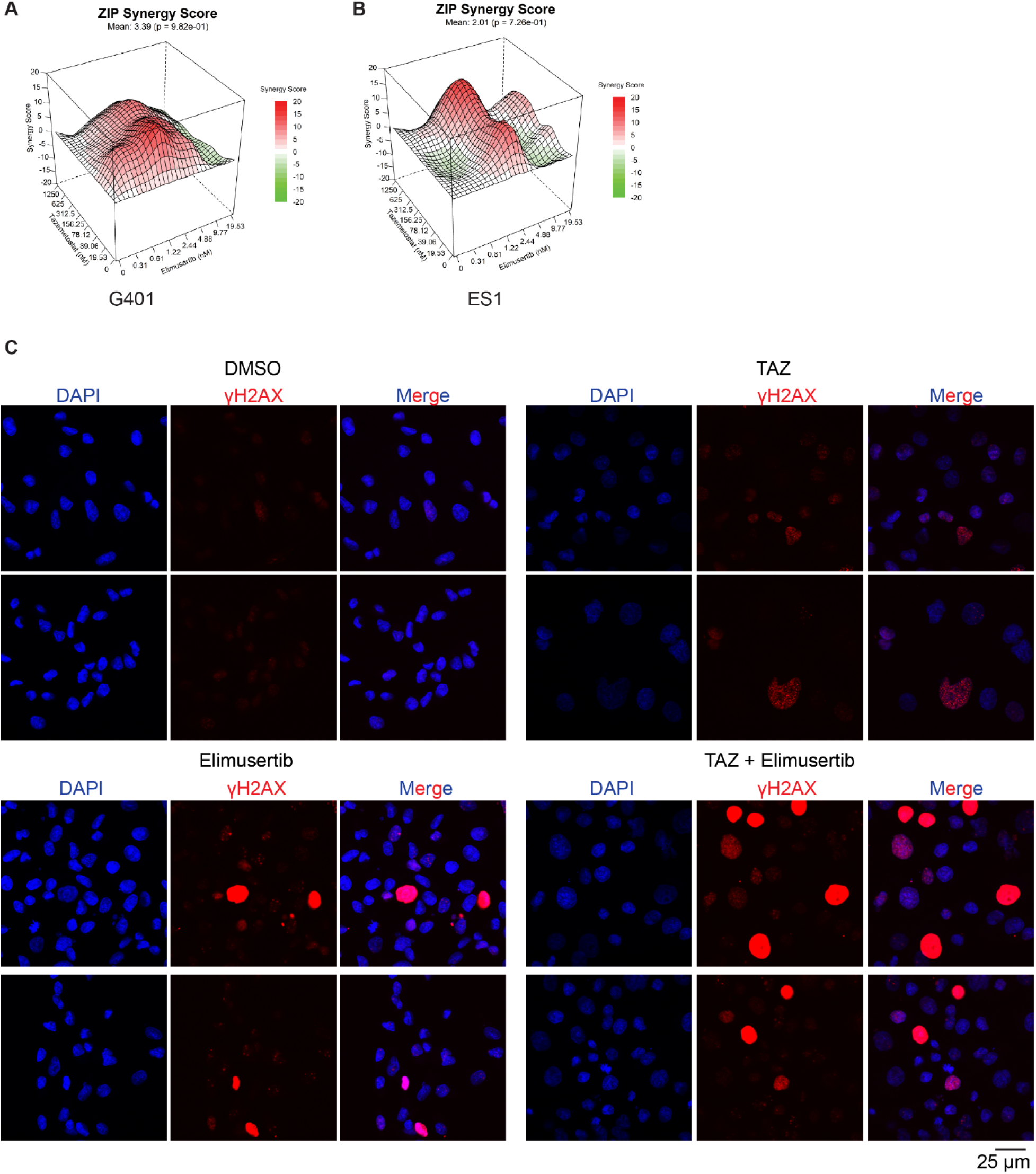
TAZ induces DNA damage in G401 cells. (**A-B**) Synergy plots for combination treatment with TAZ and elimusertib for (**A**) G401 and (**B**) ES1 cells. Cells were treated at the indicated doses for 9 days and analyzed for synergy using the Zero Interaction Potency (ZIP) m**odel.** (**C**) Representative uncropped images of G401 cells treated as indicated (same treatment as in Figure 5D). 2 fields are shown per condition.

**Figure S13:**
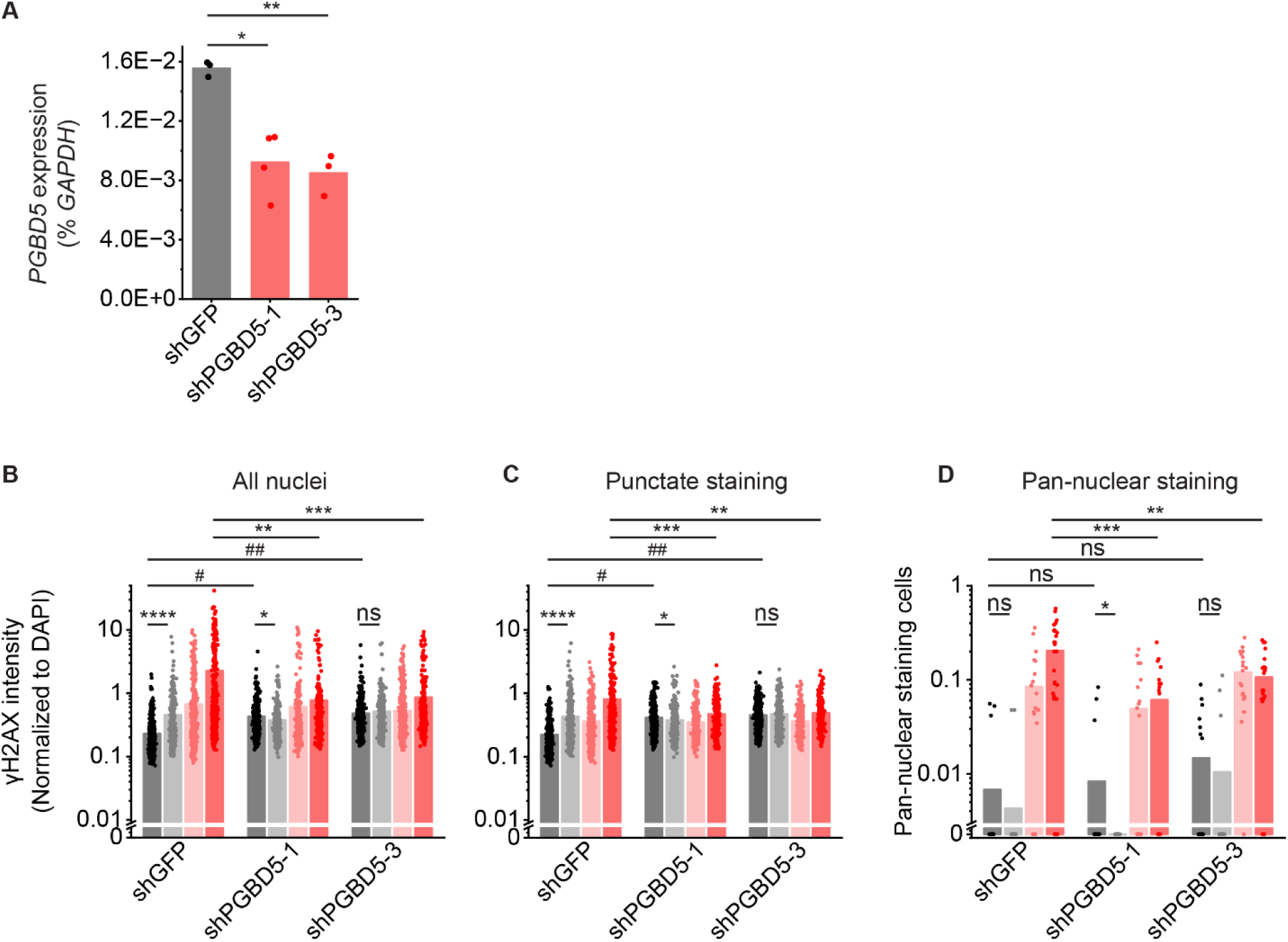
TAZ-induced DNA damage is PGBD5-dependent: (**A**) RT-qPCR showing *PGBD5* expression vs. *GAPDH* in G401 cells with the indicated shRNA. \**p* = 1.6E-2, \*\**p* = 3.8E-3. (**B**) Quantification of γH2AX fluorescence relative to DAPI fluorescence in all nuclei. \**p* = 8.6E-3, \*\**p* = 3.9E-8, \*\*\**p* = 4.4E-8, \*\*\*\**p* = 1.1E-12, # *p* = 4.9E-34, ## *p* = 4.0E-47 by two-sided Student’s t-test. (**C**) Quantification of γH2AX fluorescence relative to DAPI fluorescence in nuclei with punctate γH2AX staining. \**p* = 3.6E-2, \*\**p* = 2.7E-5, \*\*\**p* = 2.4E-5, \*\*\*\**p* = 1.1E-15, #*p* = 5.6E-51, ##*p* = 1.7E-81. For B-C, n for shGFP cells is 539 for DMSO, 321 for TAZ, 491 for elimusertib, 418 for TAZ + elimusertib. n for shPGBD5-1 is 497 for DMSO, 390 for TAZ, 354 for elimusertib, 277 for TAZ + elimusertib. n for shPGBD5-3 is 703 for DMSO, 418 for TAZ, 554 for elimusertib, 317 for TAZ + elimusertib. (**D**) Proportion of nuclei with pan-nuclear γH2AX staining per field. Each dot represents one field. \**p* = 9.2E-2, \*\**p* = 1.6E-2, \*\*\**p* = 9.0E-4 n for shGFP cells is 22 fields for DMSO, 22 for TAZ, 22 for elimusertib, 28 for TAZ + elimusertib. n for shPGBD5-1 is 22 for DMSO, 22 for TAZ, 22 for elimusertib, 20 for TAZ + elimusertib. n for shPGBD5-3 is 22 for DMSO, 22 for TAZ, 20 for elimusertib, 22 for TAZ + elimusertib.

**Figure S14:**
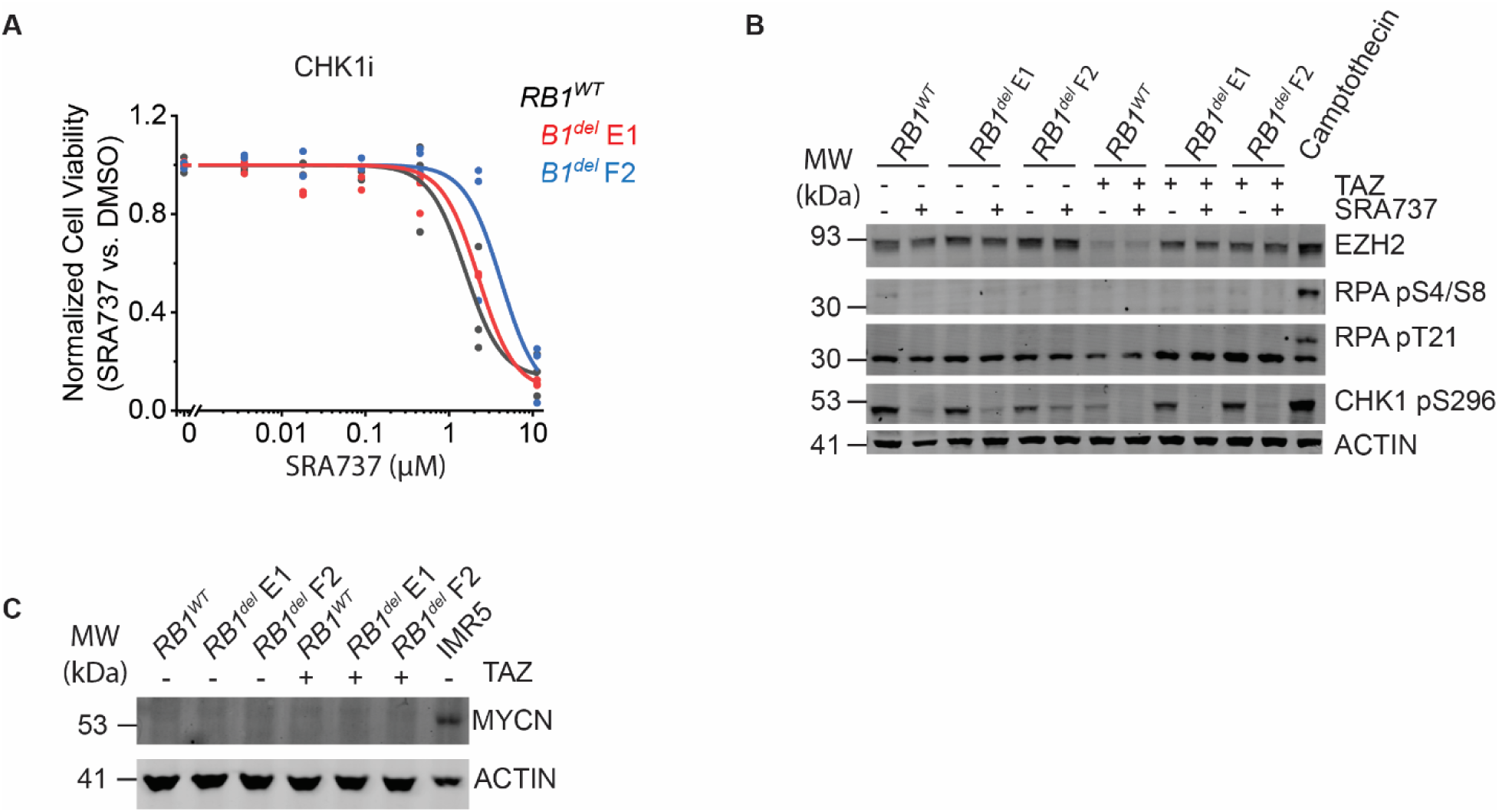
CHK1 inhibition does not induce replication stress or synergize with TAZ: (**A**) Dose-response curves of G401 cells treated with the CHK1 inhibitor SRA737 for 9 days. (**B**) Western blot assaying replication stress as measured by RPA phosphorylation at S4/8 and T21. Camptothecin treatment (1.5 µM) for 2 h was used as a positive control for replication stress. Autophosphorylation of CHK1 at S296 was used to confirm CHK1 inhibition. Cells were pre-treated with 10 µM TAZ or DMSO for 9 days. Cells were then split and additionally treated with SRA737 (3 µM) or equivalent volume of DMSO for 2 days. (**C**) Cells treated with 10 µM TAZ or DMSO for 11 days do not express MYCN protein. *MYCN*-amplified neuroblastoma cell line IMR5 was used as a positive control for MYCN expression.

**Figure S15:**
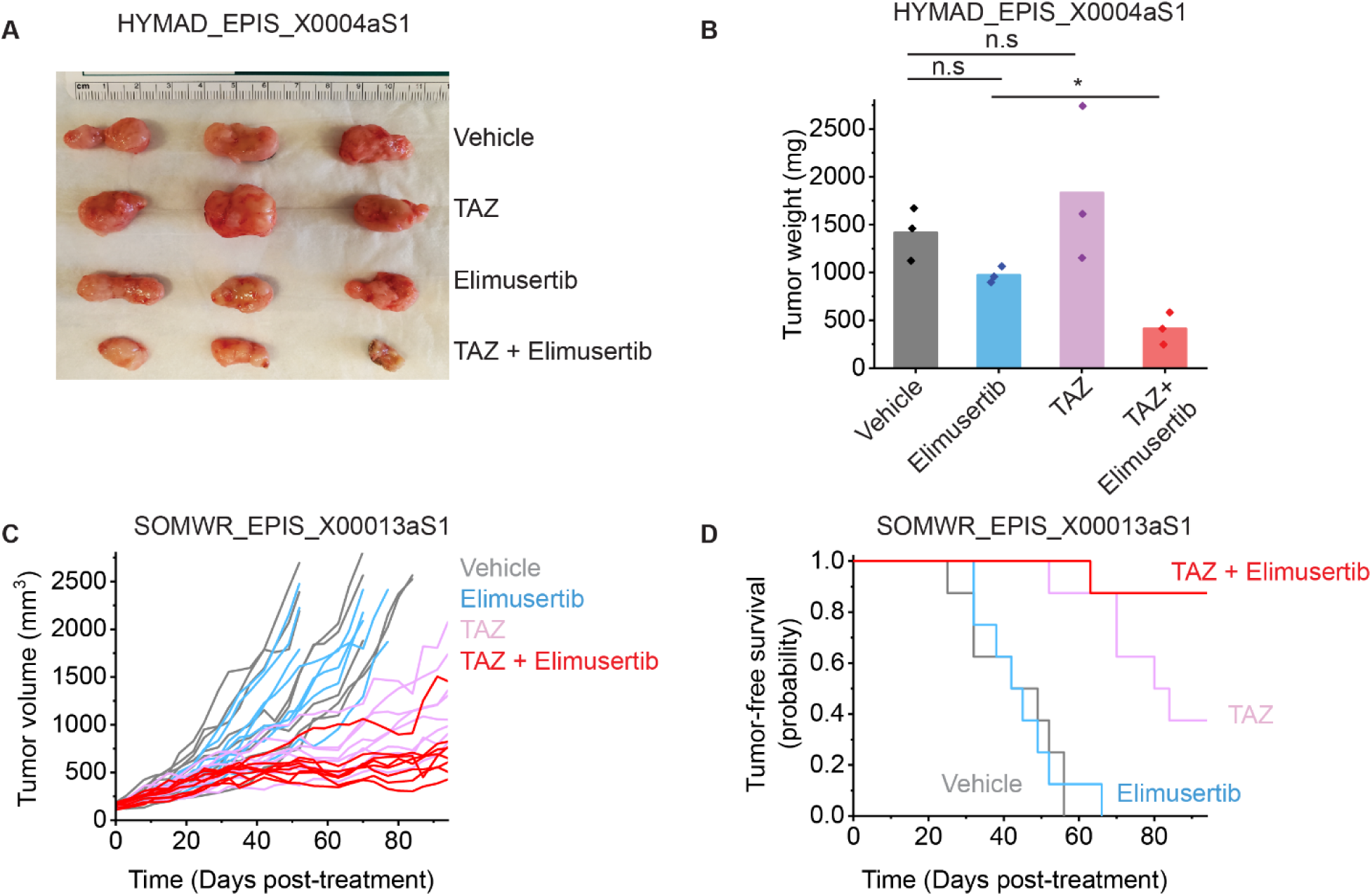
Response of individual PDX models to combination therapy: (**A**) Image of representative tumors extracted from mice in Figure 5G and on Day 52 of treatment (**B**) their corresponding weights. \**p* = 6.7E-3 by two-sided Student’s t-test. (**C**) Tumor growth curves for SOMWR_EPIS_X00013aS1 PDX model. Vardi *U*-test *p* = 1.0E-3 and 6.0E-2 for combination versus elimusertib and TAZ, respectively. *n* = 8 mice for all groups. (**D**) Kaplan-Meier curves showing tumor-free survival (defined as tumor volume ≤ 1,000 mm^3^) for SOMWR_EPIS_X00013aS1 PDX model; Mean survival is 90 days (95% CI: 83-97 days) for combination versus 80 days (95% CI: 70-90 days) for TAZ and 45 days (95% CI: 37-52 days) for elimusertib. Log-rank test *p* = 9.5E-5 and 6.0E-2 for combination versus elimusertib and TAZ, respectively.

**Figure S16:**
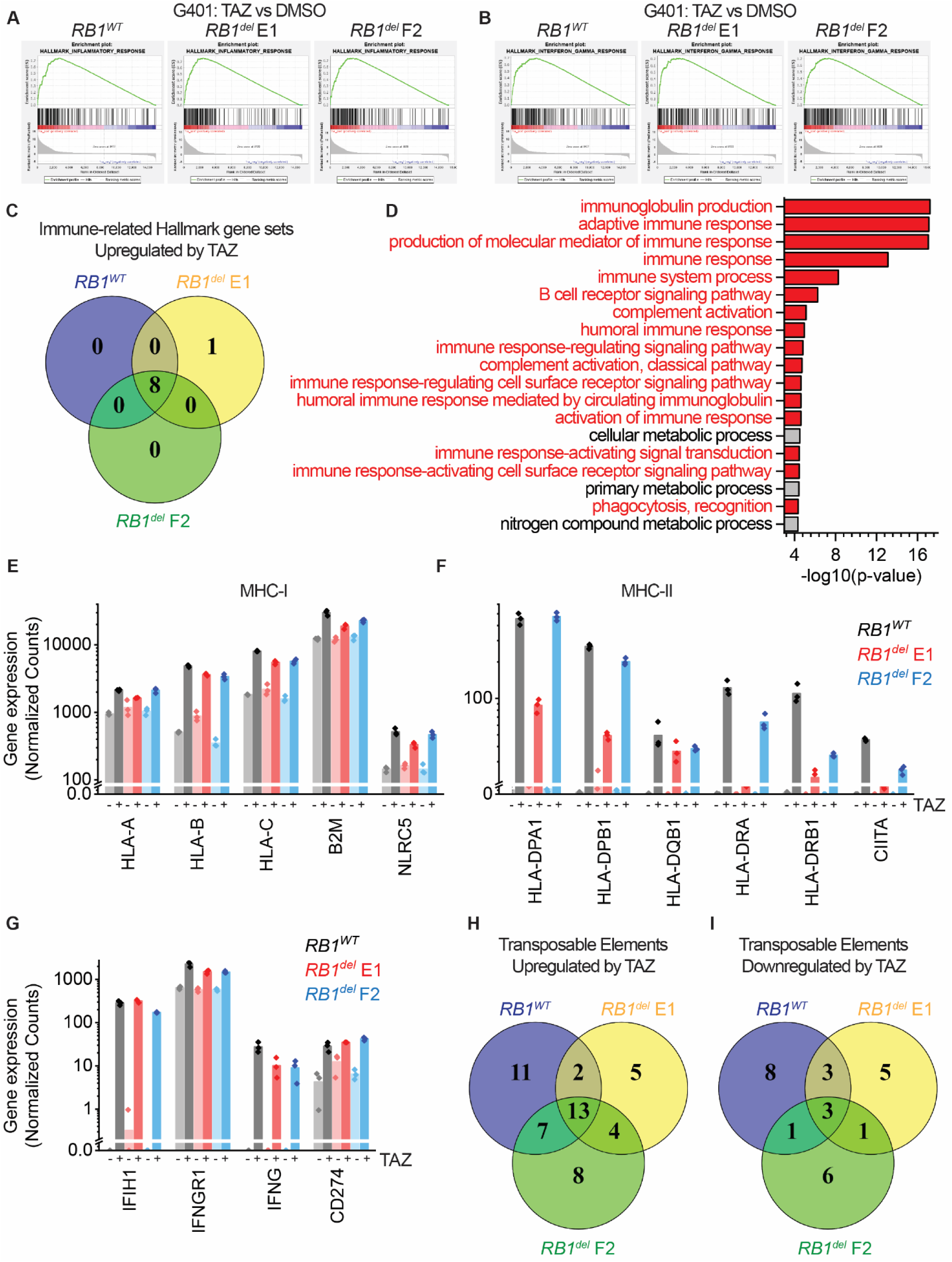
Induction of immune-related genes and endogenous transposable elements by TAZ: (**A-B**) Examples of immune-related gene sets induced by TAZ in G401 cells with and without *RB1*. Data is the same as that in Figure 2. (**C**) Venn diagrams comparing immune-related gene sets in *RB1^wt^* and *RB1^del^*cells. (**D**) Top GO terms, sorted by p-value, enriched in post-treatment patient tumor specimens that did responded to TAZ, compared to those that did not. (**E-G**) DESeq2-normalized gene expression for the indicated genes in TAZ or DMSO-treated G401 cells. Data is the same as that shown in Figure 2. (**H**) Venn diagrams comparing endogenous transposable elements up- and down-regulated by TAZ in G401 cells.

**Figure S17:**
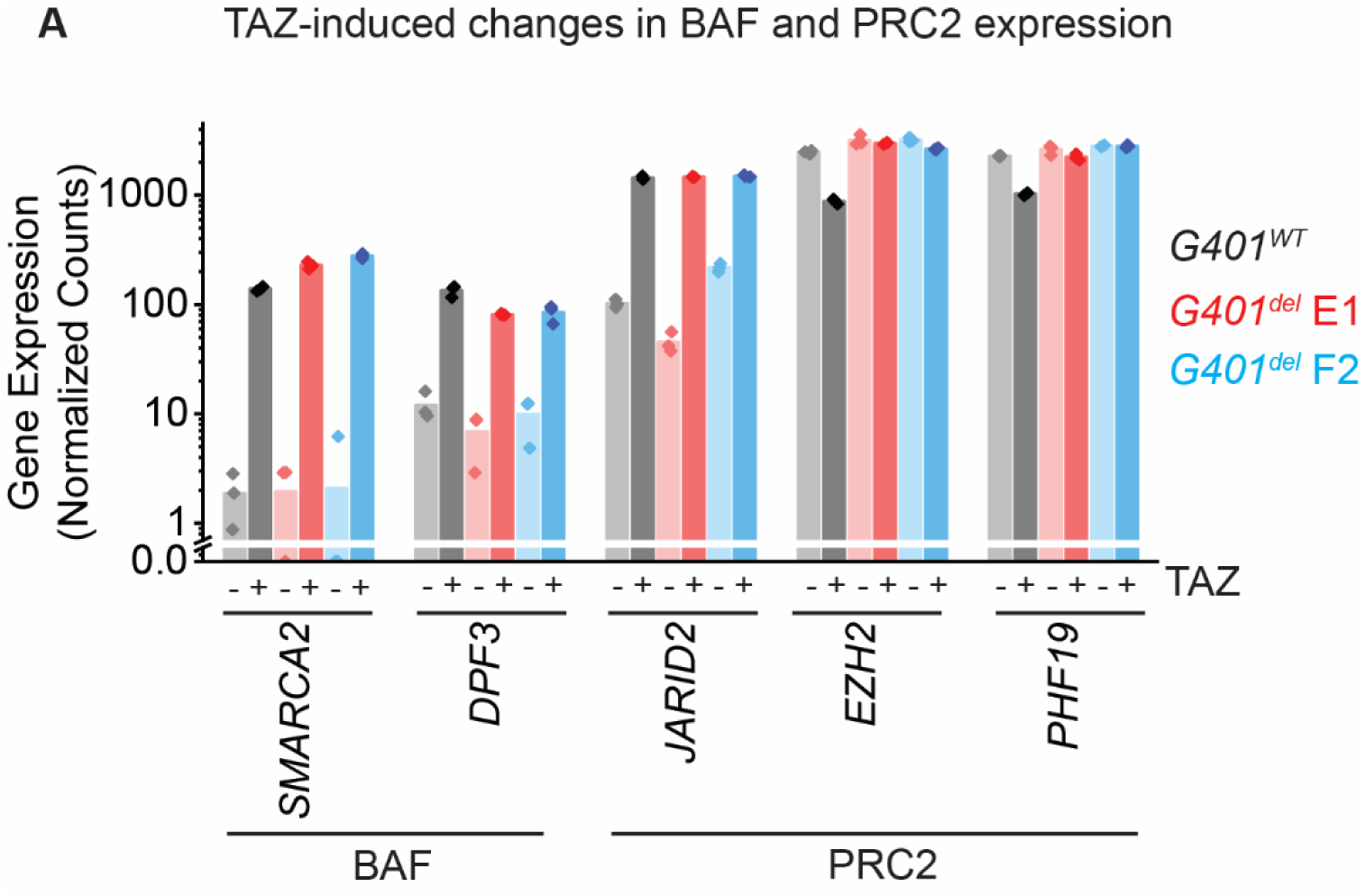
TAZ may remodel BAF and PRC2 composition by transcriptional regulation of their subunits: (**A**) DESeq2-normalized read counts for all BAF and PRC2 subunits showing significantly altered gene expression between TAZ and DMSO-treated cells. Same data as in Figure 2.

