## Supplementary material for "Overcoming clinical resistance to EZH2 inhibition using rational epigenetic combination therapy": Uncropped Western Blot

Figure 1D

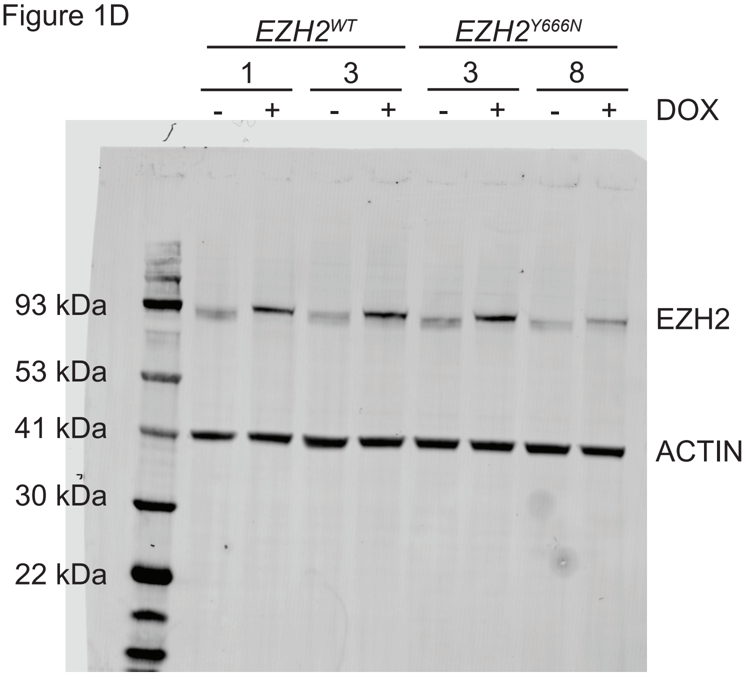


Figure 1E

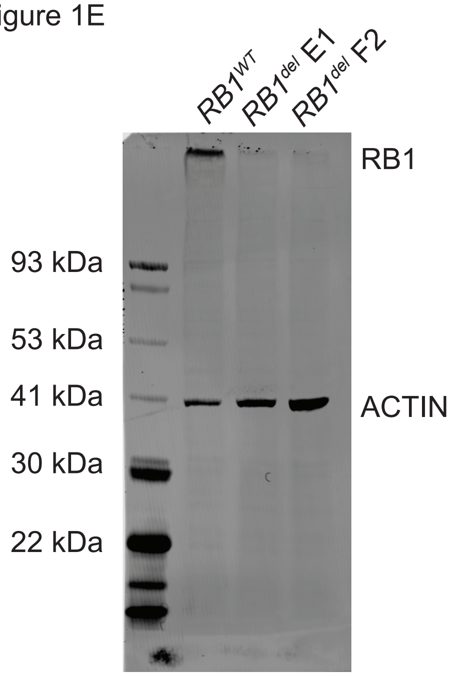


Figure 2H

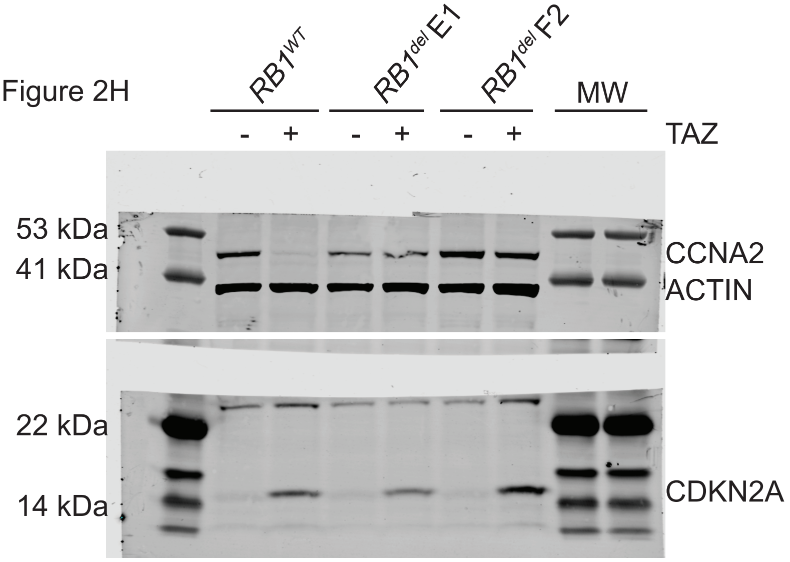


Figure 3E

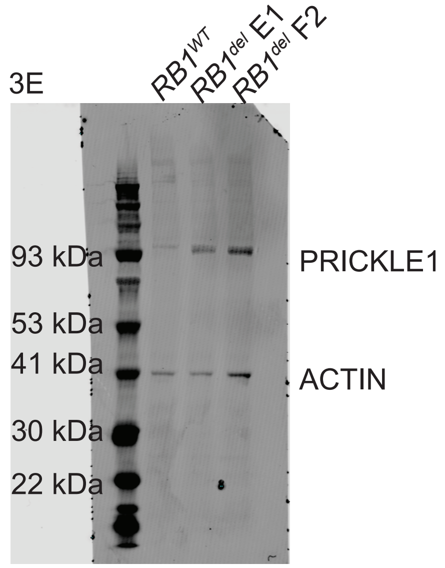


Figure S2B

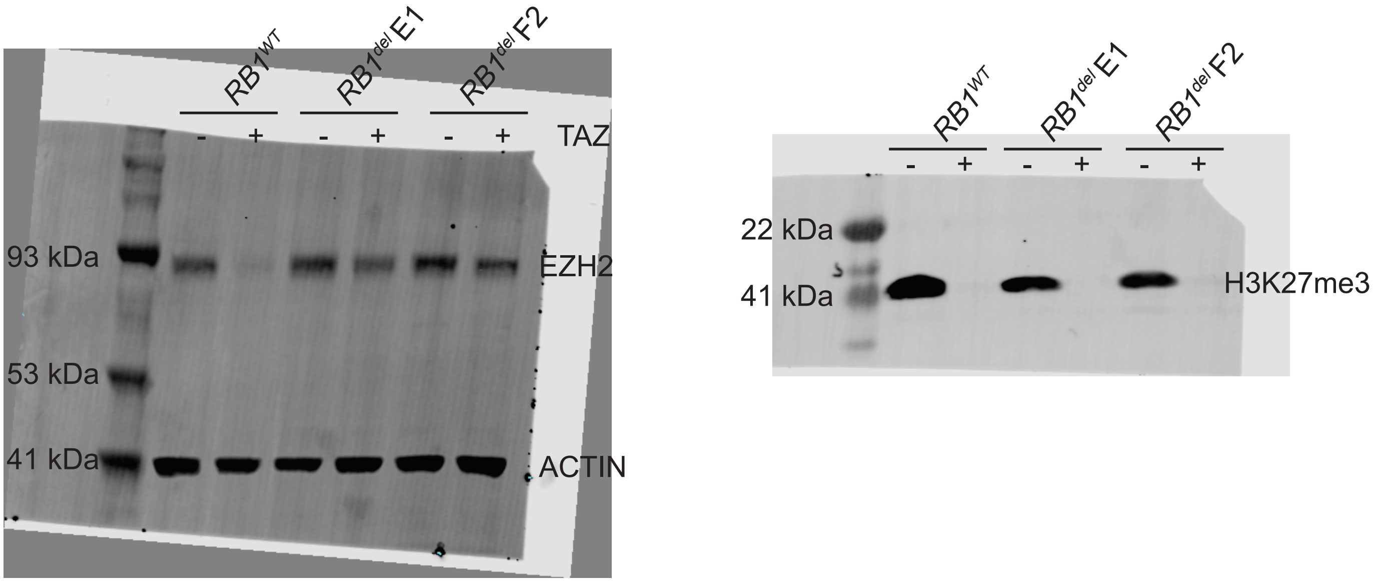

Figure S3C

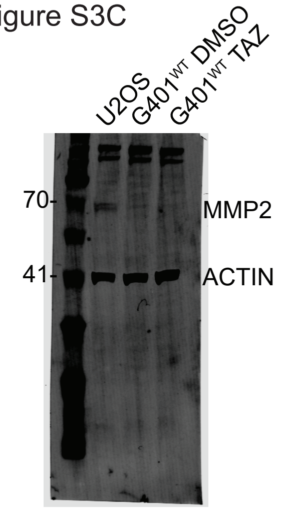


Figure S5A

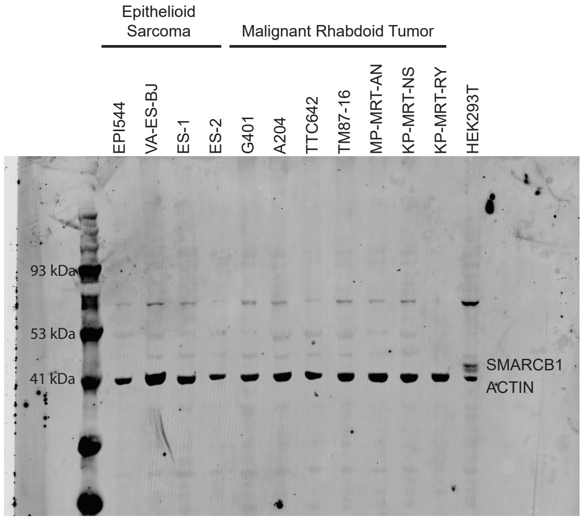


Figure S6A

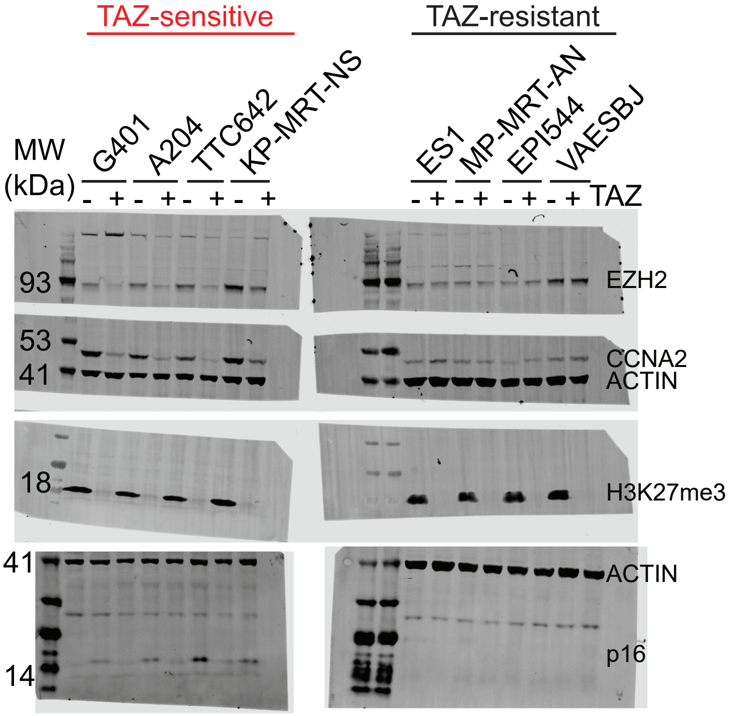

Figure S6D

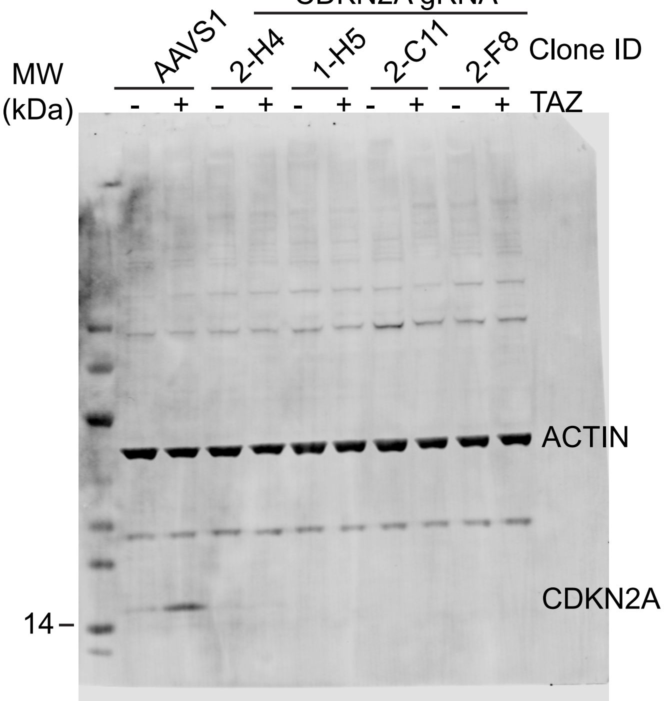


Figure S10B

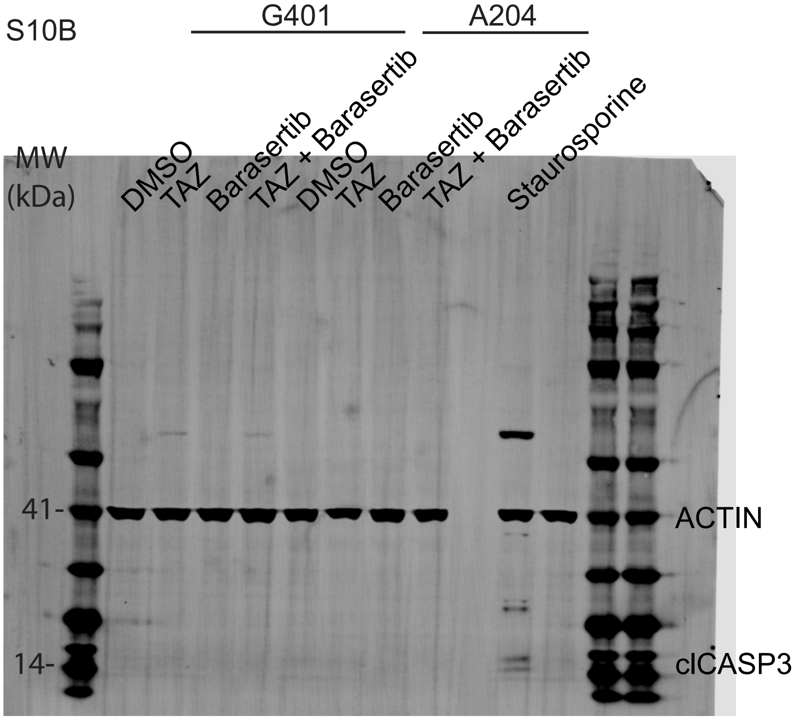


Figure S14B

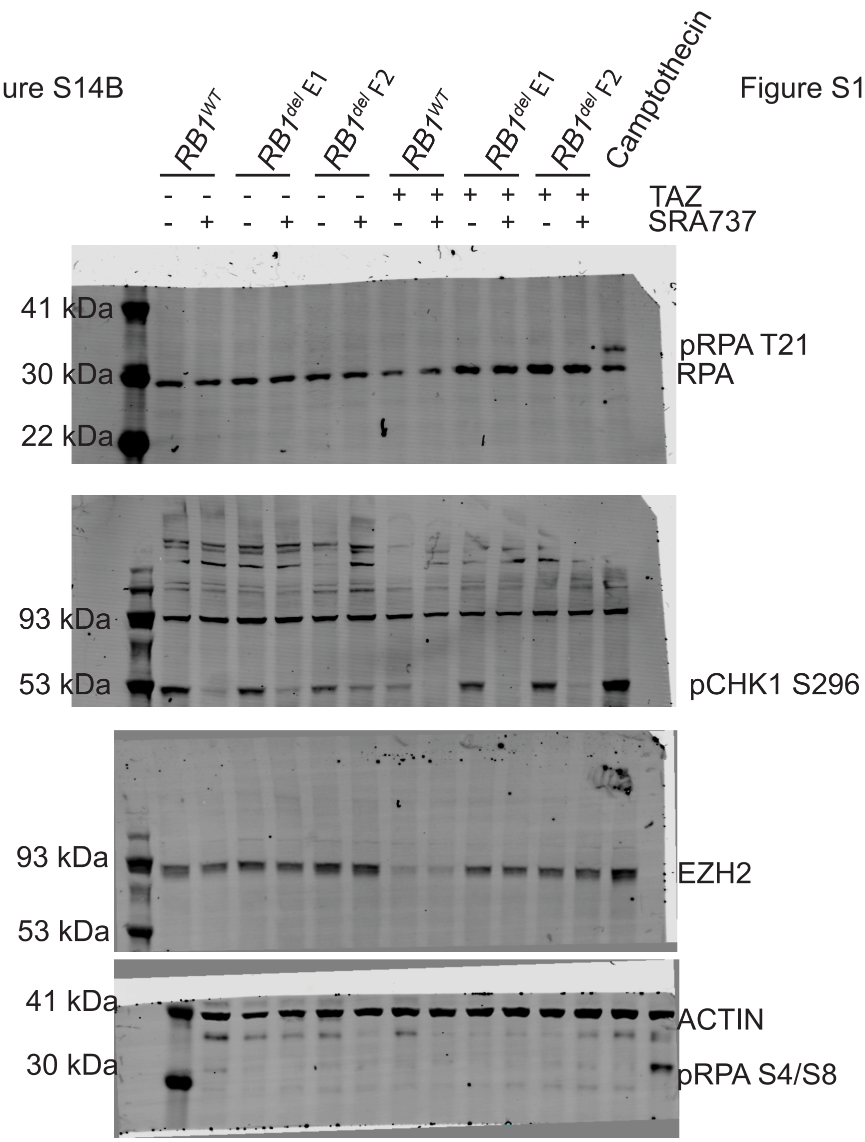


Figure S14C

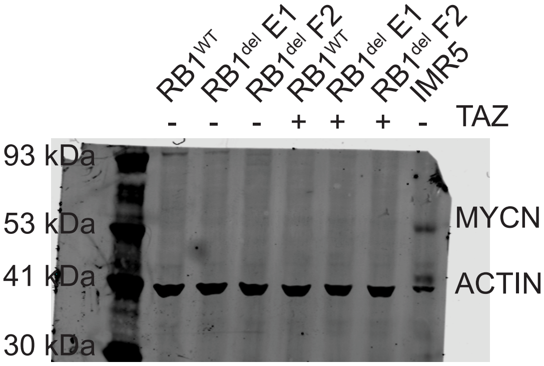
